# Single-cell analysis of pediatric acute myeloid leukemia samples uncovers treatment-resistant stem and mast cells

**DOI:** 10.1101/2024.07.09.602801

**Authors:** Denis Ohlstrom, Mojtaba Bakhtia, Hope Mumme, Marina Michaud, Frank Chien, William Pilcher, Sarthak Satpathy, Sean Jordan, Swati Bhasin, Manoj Bhasin

**Author notes:** Corresponding author: Manoj K. Bhasin, MS, PhD, Aflac Cancer and Blood Disorders Center, Children Healthcare of Atlanta, Health Sciences Research Building II, Room N320, 1760 Haygood Drive, 3^rd^ Floor, Emory School of Medicine, Atlanta, GA 30322.

## Abstract

Pediatric acute myeloid leukemia (pAML) is a heterogeneous malignancy driven by diverse cytogenetic mutations. While risk stratification improved by identifying cytogenetic lesions, prognostication remains inadequate with 30% of standard-risk patients experiencing relapse within 5 years. Single-cell RNA sequencing (scRNAseq) enabled the interrogation of malignant cell heterogeneity in pAML and characterization of the immune microenvironment. Herein we report the largest pAML scRNAseq analysis to date with 708,285 cells from 164 bone marrow biopsies of 95 patients and 11 healthy controls. We uncovered treatment-resistant (TR) subtypes of pAML specific to RUNX1-RUNX1T1, FLT3-ITD, and CBFB-MYH11 patients. The enrichment of TR subtype gene signatures on the TARGET pAML data supported an association with significantly poor outcomes. Intriguingly, in addition to leukemic stem cells, we identified mast cell-like pAML associated with treatment resistance and poor outcomes. Together, immature and mature pAML subtypes are promising biomarkers for identifying patients at increased risk of relapse within cytogenetic categories.

## Introduction

Pediatric acute myeloid leukemia (pAML) is a heterogeneous malignancy caused by diverse driver mutations. Common cytogenetic aberrations include internal tandem duplications of FLT3 (FLT3-ITD, ∼20%) or translocations in KMT2A/MLL (∼20%), RUNX1-RUNXT1T1 (15%), and CBFB-MYH11 (10%)^1,2^. While the identification of driver lesions has improved patient risk stratification, standard-risk lesions still have 5-year relapse rates as high as 30%^3–5^. Thus, biomarkers that differentiate early relapsing pAML within existing cytogenetic categories are needed.

Several recent single-cell RNA sequencing (scRNAseq) studies characterized the diverse composition of malignant cell types in pAML, including those resembling monocytes, granulocyte-monocyte progenitors (GMPs), dendritic cells (DCs), and hematopoietic stem cells (HSCs) among others^6–9^. Our lab recently profiled pAML samples at diagnosis and end-of-induction, identifying a risk-associated gene set for treatment-resistant blast cells^9^. Leukemic stem cells (LSCs) are a CD34^+^ subtype of pAML (analogous to the HSC) that persist through treatment and are also considered responsible for relapse^10,11^. Recent scRNAseq studies have proposed that LSCs express unique cell surface markers such as CD69 and are associated with poor outcomes^12^. Conversely, mast cells are well-differentiated myeloid cells that typically comprise less than 1% of the bone marrow (BM) microenvironment and are frequently undetected or not annotated in scRNAseq experiments^13^. Previous immunohistochemical studies found up to 40% of adult AML patients with core binding factor mutations (CBF, comprised of RUNX1-RUNX1T1 and CBFB-MYH11 mutations) have elevated BM mast cells^14,15^. No large studies of pAML have examined mast cells, though case reports of systemic mastocytosis with associated hematological neoplasm (SM-AHN) have described increased mast cells in patients with co-occurring CBF pAML^16–22^. Large scRNAseq studies are needed to examine the heterogeneity of LSCs and well-differentiated pAML subtypes to better understand the subtypes that drive relapse within cytogenetic categories. Identifying treatment-resistant cellular subtypes could provide functional insights for selecting specifically efficacious treatments to improve remission durability.

To this end, we analyzed 708,285 cells from 164 BM biopsies, including samples from 95 patients, and 11 healthy controls to identify cell types associated with treatment resistance and poor outcomes. High-risk, treatment-resistant (HRTR) cellular subtypes were identified within cytogenetic categories based on greater abundance in patients who experienced relapse, persistence after treatment, and gene expression associated with poor outcomes in the Therapeutically Applicable Research to Generate Effective Treatments (TARGET) bulk RNAseq data. We also examined unique transcriptomic features of HRTR subtypes and their influence on the immune microenvironment. Finally, the deconvolution of the Beat-AML dataset enabled the identification of drugs or small molecular inhibitors that are specifically efficacious against HRTR subtypes with subsequent *in vitro* validation (**Fig 1**). This study represents the largest scRNAseq analysis of pAML to date. Matched diagnosis, EOI, and relapsed samples enabled the identification of multiple HRTR pAML subtypes specific to cytogenetic groups and provided preliminary insights for their treatment. The study also reveals a potent association of elevated mast cells with poor outcomes in core binding factor pAML.

**Figure 1:**
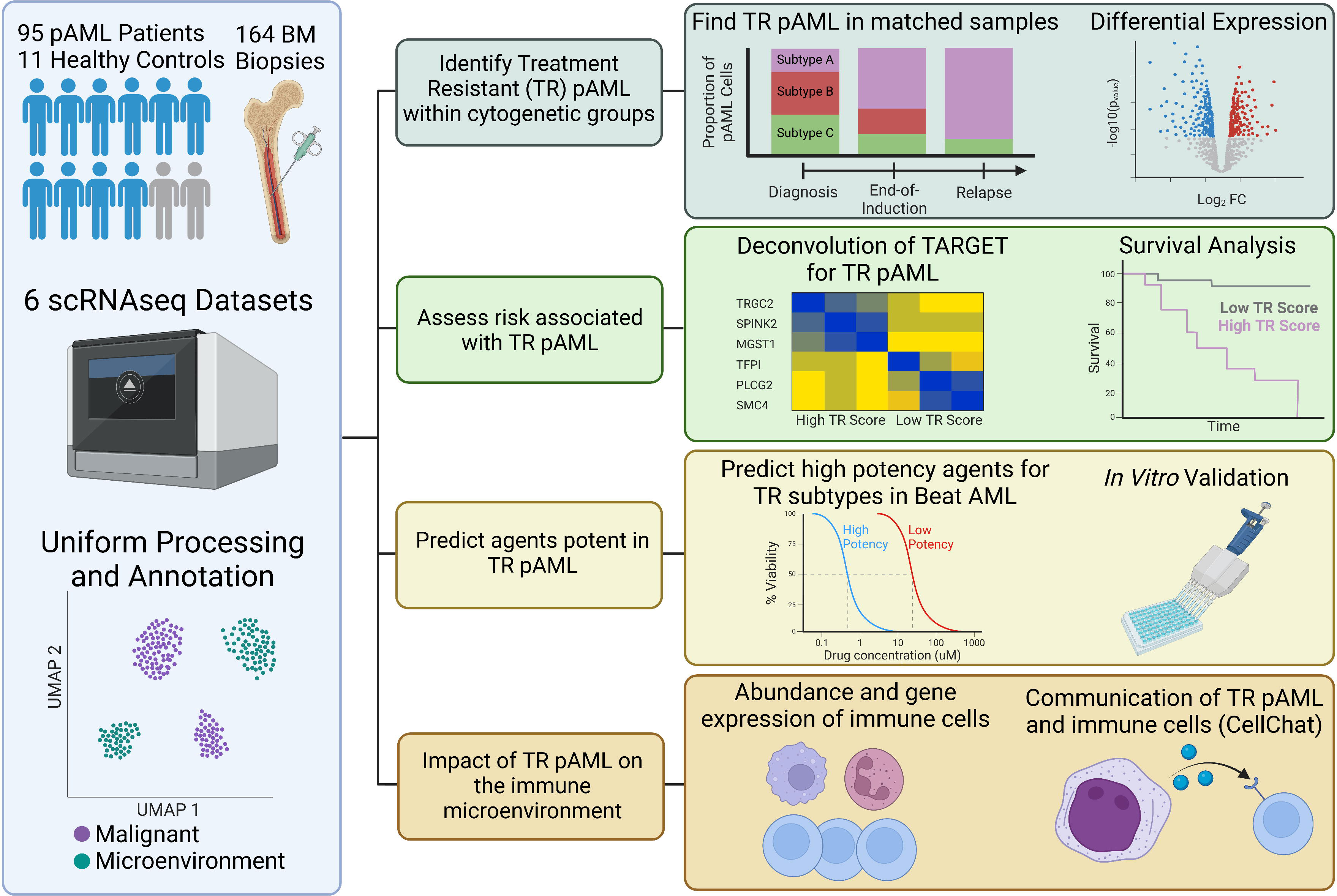
Schematic for analysis of pediatric acute myeloid leukemia datasets to identify and characterize high-risk, treatment-resistant subtypes.

## Results

### Generation of uniformly processed and annotated pAML dataset

Six scRNAseq datasets were obtained from the NCBI GEO and scPCA repositories, including two from our laboratory (**Supplemental Table 1**)^9^. The datasets included 164 BM biopsies from 95 pAML patients (ages 0.2-24.4 years) and 11 young adult healthy controls (ages 19-26, **Supplemental Table 2**). Ninety-one samples were collected at the time of disease diagnosis (DX), 34 at end-of-induction (EOI), and 28 at relapse (REL). Documented outcomes were available for 53 patients, with 22 sustained remissions and 31 relapses. Thirty-seven patients had matched samples, including 36 from DX, 33 from EOI, and 26 from REL.

To uniformly process each dataset, we removed low-quality cells (>20% mtRNA,<250 or >6000 features) and doublets (DoubletFinderV3^23^). The resulting feature-cell barcode matrices (708,285 total cells) were annotated using canonical cell type markers and integrated using the Seurat integration anchors approach^24^ (**Fig 2A-B**, **Supplemental Table 3**). The proportion of cell types was comparable across datasets (p=0.944, **Fig 2C**).

**Figure 2:**
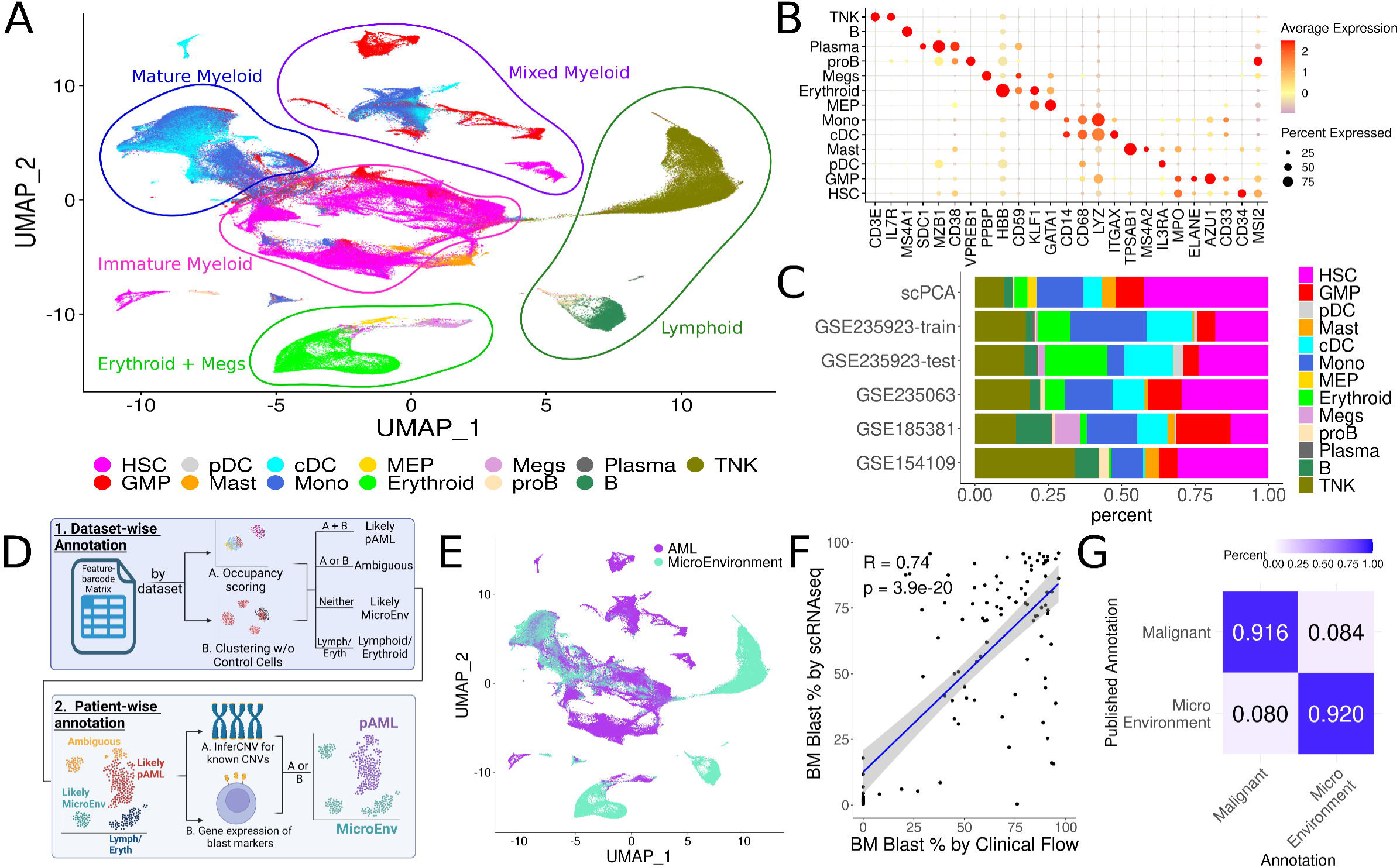
Malignant blast cell identification. (A) UMAP embedding of 164 pAML samples derived from 95 patients and 11 healthy controls. 708,285 cells passed quality control and were annotated by expression of canonical cell type gene markers. (B) Dot plot showing expression of canonical cell type gene markers. The color intensity represents gene expression (red: high, yellow: medium, blue: low) and the size of dots represents the percent of cells expressing the marker gene. (C) Bar chart showing the relative proportion of annotated cell types in each dataset. The proportion of cells was not found to be different between datasets by two-way ANOVA (p = 0.944). (D) Schematic for annotation of malignant myeloid cells. (E) UMAP embeddings of all cells colored by malignant (purple) or microenvironment (turquoise) annotation. (F) Scatter plot of bone marrow blast percentage by flow cytometry vs percent of cells annotated as malignant in scRNAseq. The relationship between percentages was assessed by Pearson correlation coefficient testing. (G) Confusion matrix for consistency between malignant/microenvironment annotations produced by the method described herein versus annotations published with GSE185381. The color of each box indicates the proportion, with blue for a proportion approaching 1.0 and white for a proportion approaching 0.0. UMAP: Uniform manifold approximation and projection, scRNAseq: single-cell RNA sequencing, HSC: hematopoietic stem cell, GMP: granulocyte monocyte progenitor, cDC: classical dendritic cell, Mono: monocyte, MEP: monocyte erythrocyte progenitor, RBC: red blood cell, Megs: Megakaryocytes, CNV: copy number variant, BM: bone marrow.

### Multiparameter annotation of malignant pAML cells

To robustly annotate myeloid clusters as malignant or non-malignant, microenvironment, we applied a two-step annotation approach (**Fig 2D**). First, cells were clustered dataset-wise using the Louvain algorithm, yielding 412 clusters (an average of 1,719 cells per cluster) across 6 datasets. For each cluster, we calculated three metrics to aid in discriminating malignant versus microenvironment clusters: 1.a) The percentage of cells derived from a single patient (patient-specific occupancy scoring as described previously^6^), 1.b) The percentage of cells derived from patients with the same cytogenetic lesion, and 2) The percentage of cells from healthy control samples (**Supplemental Table 4**). First, a cluster was labeled “likely malignant” if it contained<2% healthy control cells and >70% of cells specific to a single patient or cytogenetic group (**Supplemental Figure 1-2**). Second, cells were analyzed patient-wise for expression of malignant blast markers and the presence of clinically detected copy number variants (CNVs) predicted using the inferCNV algorithm^25^ (**Supplemental Figure 3**). Cells with blast marker expression or CNV detection had their annotation updated to malignant (“AML”), while cells negative for both were annotated as microenvironment (**Fig 2E, Supplemental Figure 4**). The resulting annotations labeled 53.08% of cells as malignant, a proportion highly correlated to blast cell percentage reported by clinical-standard flow cytometry (R=0.74, p<0.05, **Fig 2F**). Furthermore, our malignant and microenvironment annotations were 91.6% and 92.0% consistent, respectively, with published annotations for the GSE185381 dataset (**Fig 2G**). Thus, our multiparameter approach leveraged dataset-wise and patient-wise features to robustly identify malignant cells across all datasets.

### Analysis of matched diagnosis, end-of-induction, and relapse samples reveals treatment-resistant pAML cell subtypes

To identify treatment-resistant pAML blast cell subtypes we examined matched DX, EOI, and REL samples from 37 patients. These samples included seven RUNX1-RUNX1T1, four CBFB-MYH11, five FLT3-ITD, and twelve KMT2A rearranged samples (referred to as RUNX1, CBFB, FLT3, and KMT2A respectively in the manuscript) and 9 patients with various alterations grouped as “other” (**Fig 3A-B**, **Supplemental Table 2**). Malignant blast cells from matched samples were merged, batch-corrected, and clustered using the Louvain algorithm. The resulting 60 clusters were annotated into 33 subtypes based on similar top differentially expressed genes corresponding to cell surface proteins (Log_2_FC > 0.5, adjusted p<0.05, **Fig 3C**). Of the 33 pAML subtypes, 11 were HSC-like (*CD34*^+^), eight GMP-like (*AZU1*^+^, *ELANE*^+^), eight monocyte-like (*CD14*^+^, *CD68*^+^, *LYZ*^+^), three cDC-like (*ITGAX*^+^), two mast-like (*TPSAB1*^+^) and one megakaryocyte-like (*PPBP*^+^) (**Fig 3D**). Consistent with the diversity of KMT2A rearrangements^26^ and “other” categories, 17 out of 21 patients formed patient-specific subtypes (>90% of cells in a subtype from one patient, **Supplemental Figure 5**). RUNX1, FLT3, and CBFB pAML subtypes overlapped between patients with only two out of sixteen patients forming patient-specific subtypes (**Supplemental Figure 5**). Therefore, we focused our further analysis on identifying high-risk treatment-resistant (HRTR) pAML subtypes within the RUNX1, FLT3, and CBFB cytogenetic groups. Three criteria defined HRTR pAML subtypes: 1) Proportionally greater representation in patients who experienced relapse versus sustained remission, 2) Proportional increase in cell-type abundance at EOI versus DX, and 3) Overexpression of a gene signature associated with poor outcomes.

**Figure 3:**
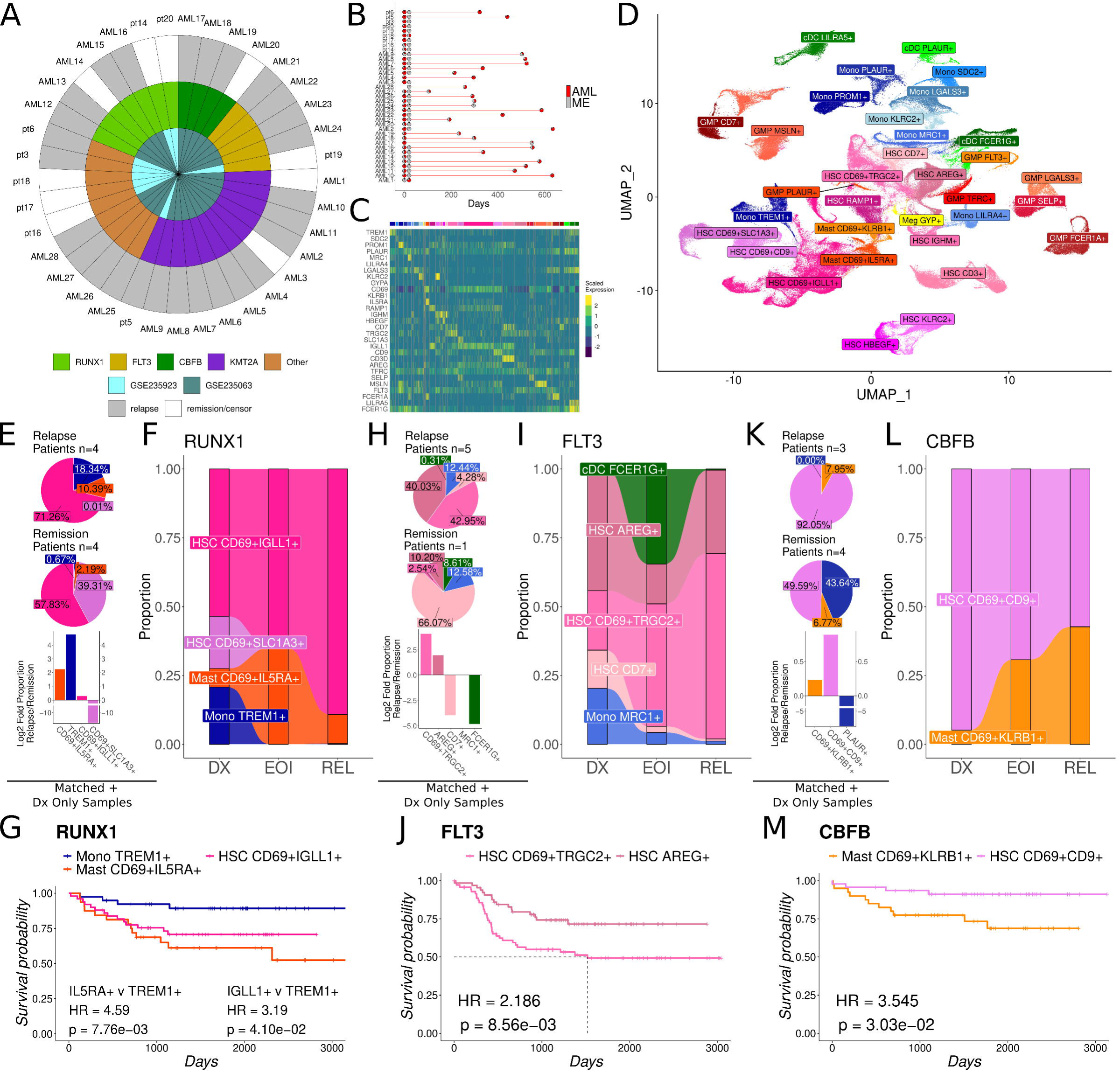
Analysis of malignant cells from matched diagnosis, end-of-induction, and relapse samples reveals treatment-resistant cell types. (A) Donut plot showing the source dataset (inner), cytogenetic group (middle), and relapse/remission outcome for 37 patients with matched sample. (B) Scatterpie chart for each patient with matched samples. Each patient is represented by pie charts at the same y-axis level. Pie charts indicate the proportion of malignant cells (red) or microenvironment cells (gray), positioned along the x-axis at the number of days after diagnosis the biopsy was taken. (C) Heatmap of the 30 genes coding for cell surface markers identified as over-expressed (log_2_ fold change > 0.5, adjusted p-value < 0.05, Wilcoxon rank sum test) in pAML subtypes. The Y-axis represents gene names and the X-axis is grouped and colored by the pAML subtypes. Color scale: yellow indicates higher expression, blue indicates lower expression. (D) UMAP embeddings of malignant cells colored and labeled by myeloid cell type and over-expressed cell surface genes. (E) Pie charts showing the percentage of cells from patients who experienced relapse (top) or sustained remission (middle) that were annotated into different pAML subtypes. Bar chart of the log_2_ fold proportion of cells (relapse divided by remission) (bottom). Positive values indicate relapse-associated subtypes. Color indicates pAML subtypes as in D and F. (F) Alluvial plot of the proportion of cells in each pAML subtype from RUNX1 patients at diagnosis (DX), end-of-induction (EOI), and relapse (REL). Subtypes that were proportionally increased at EOI vs DX were considered treatment-resistant. (G) Kaplan Meier curve for overall survival of samples from the TARGET database with similar gene expression to pAML subtypes in scRNAseq. Samples were considered enriched for a subtype based on high expression of pAML subtype gene signatures (see Supplemental Figure 6). Cox proportional hazard models were used for the calculation of hazard ratio and significance testing. (H-J) Plots as in E-G for FLT3 pAML subtypes. (K-M) Plots as in E-G for CBFB pAML subtypes. pAML: pediatric acute myeloid leukemia, ME: microenvironment, HSC: hematopoietic stem cell, GMP: granulocyte-monocyte progenitor, cDC: classical dendritic cell, Mono: monocyte, Meg: megakaryocyte, HR: hazard ratio.

For RUNX1 pAML, three subtypes met the first HRTR criteria (enrichment in patients who experienced relapse): *CD69*^+^*IL5RA*^+^ mast-like pAML (10.39% relapse vs 2.19% remission), *CD69*^+^*IGLL1*^+^ HSC-like pAML (71.26% relapse vs 57.83% remission), and *TREM1*^+^ monocyte like pAML (18.34% relapse vs 0.67% remission, **Fig 3E**). Of these, two subtypes met the second HRTR criteria (proportional increase at EOI versus DX): *CD69*^+^*IGLL1*^+^ HSC-like pAML (DX=53.5%, EOI=65.0%) and *CD69*^+^*IL5RA*^+^ mast-like pAML (DX=6.8%, EOI=35.0%), while *TREM1*^+^ monocyte-like pAML was depleted after treatment (DX=20.8%, EOI=0.0%, **Fig 3F**). For the third criterion, we evaluated overall survival for samples in the TARGET pAML dataset with similar gene expression to relapse-associated pAML subtypes. Gene signatures for relapse-associated subtypes contained the top 30 highly overexpressed genes if a large number of genes achieved the significant dysregulation threshold (Log_2_FC > 0.5 and adjusted p<0.05). Diagnostic BM samples in the TARGET pAML dataset from patients with RUNX1-RUNX1T1 mutation were scored for the three relapse-associated gene signatures by average normalized, z-scaled expression. TARGET pAML samples were labeled “enriched” for the highest subtype signature score (e.g. *CD69*^+^*IGLL1*^+^ HSC-like enriched if that was the highest signature score) or “none” if all signature scores were negative (**Supplemental Figure 6**). In RUNX1 pAML, enrichment for the treatment-resistant *CD69*^+^*IGLL1*^+^ HSC-like (HR=3.19, p<0.05) and *CD69*^+^*IL5RA*^+^ mast-like (HR=4.59, p<0.05) signatures were associated with worse outcomes compared to enrichment for the *TREM1*^+^ monocyte-like signature (**Fig 3G**).

In FLT3 pAML, two subtypes met the first HRTR criteria (relapse enrichment): *CD69*^+^*TRGC2*^+^ HSC-like pAML (42.95% relapse, 2.54% remission) and *AREG*^+^ HSC-like pAML (40.03% relapse, 10.20% remission **Fig 3H**). *CD69*^+^*TRGC2*^+^ HSC-like pAML met the second HRTR criteria, proportional increase at EOI (Dx=21.6%, EOI=44.5%), while *AREG*^+^ HSC-like pAML was proportionally decreased (DX=44.0%, EOI=14.4%, **Fig 3I**). For the third criterion (poor outcome association), deconvolution of FLT3 samples in TARGET (**Supplemental Figure 6**) found the *CD69*^+^*TRGC2*^+^ HSC-like gene signature was associated with poor outcomes compared to the *AREG*^+^ HSC-like signature (HR=2.19, p<0.05, **Fig 3J**).

In CBFB pAML, two subtypes were proportionally enriched in patients who experienced relapse: *CD69*^+^*KLRB1*^+^ mast-like pAML (7.95% relapse vs 6.77% remission) and *CD69*^+^*CD9*^+^ HSC-like pAML (92.05% relapse vs 48.27%, **Fig 3K**). *CD69*^+^*KLRB1*^+^ mast-like pAML was enriched at EOI (DX=5.1%, EOI=30.8%, **Fig 3K**), while *CD69*^+^*CD9*^+^ HSC-like pAML was proportionally decreased after treatment (DX=94.5%, EOI=69.2%, **Fig 3L**). Enrichment for the *CD69*^+^*KLRB1*^+^ mast-like gene signature in CBFB samples from the TARGET AML dataset was associated with poor overall survival compared to the *CD69*^+^*CD9*^+^ HSC-like gene-signature (HR=3.55, p<0.05, **Fig 3M**). Cumulatively these analyses uncovered immature and mature HRTR subtypes of pAML that are over-represented in patients who experienced relapse, persist through treatment, and express gene signatures associated with poor overall survival in the TARGET AML dataset.

### High Risk, Treatment-resistant CD69^+^IGLL1^+^ HSC cells are proliferative, anti-apoptotic, and immune effectors

To further understand the molecular landscape that contributes to poor outcomes associated with HRTR RUNX1 pAML, we examined the marker genes expressed by *CD69*^+^*IGLL1*^+^ HSC-like cells (**Fig 4A**). *CD69*^+^*IGLL1*^+^ HSC-like pAML cells significantly overexpressed 109 genes and downregulated 152 genes with absolute Log_2_FC (L_2_FC) > 0.25 and adjusted p<0.05 as compared to other AML cell types (**Fig 4B**, **Supplemental Table 6**). Pathway analysis identified a significant association with proliferative, anti-apoptotic, and immune interaction pathways (**Fig 4C**)^27^. *CD69*^+^*IGLL1*^+^ HSC-like pAML highly expressed genes known to be risk-associated (*EGFL7* L_2_FC=1.10, *BAALC* L_2_FC=0.81, *STAT1* L_2_FC=0.51, adjusted p<0.05 for all)^28–31^, immune effectors (*C1QTNF* L_2_FC=1.28, *PRTN3* L_2_FC=1.64, *CYTL1* L_2_FC=1.03, adjusted p<0.05 for all)^32,33^, and pro-proliferative mediators such as *EGFL7*^34^. To examine the association of *CD69*^+^*IGLL1*^+^ HSC-like pAML with proliferation, we compared its gene signature score with clinical BM blast percentage in the TARGET AML data and observed a significant positive correlation (R=0.26, p<0.05, **Fig 4E**). Thus, DE and pathway analysis indicated that *CD69*^+^*IGLL1*^+^ HSC-like pAML promotes poor outcomes via proliferative and anti-apoptotic gene expression.

**Figure 4:**
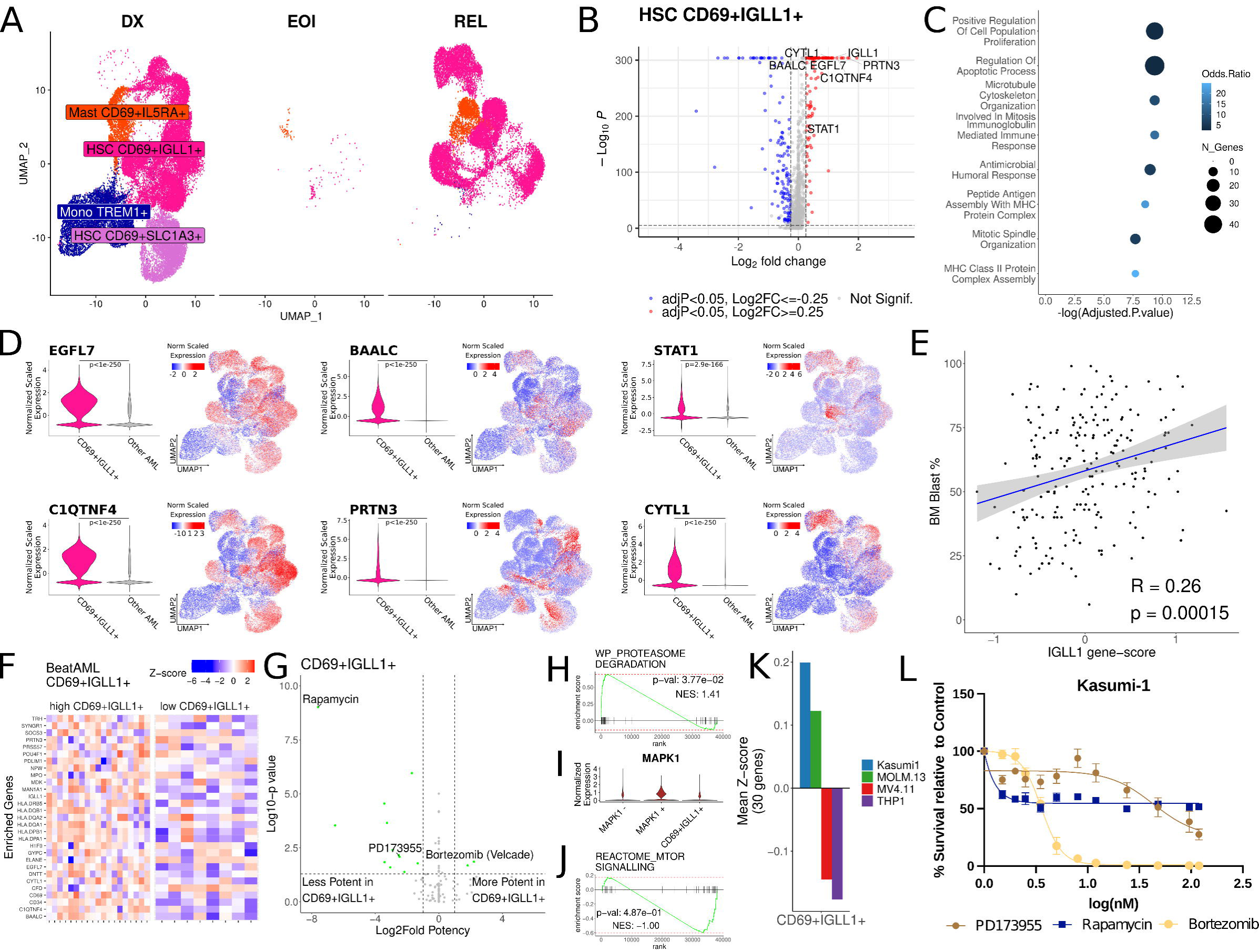
High-risk, treatment-resistant cells in RUNX1 pediatric acute myeloid leukemia (pAML) are proliferative, anti-apoptotic, and immune effectors. (A) UMAP embeddings depicting the RUNX1 pAML cells at diagnosis (DX) end-of-induction (EOI) and relapse (REL). (B) Volcano plot of differentially expressed genes in *CD69*^+^*IGLL1*^+^ HSC-like pAML vs all other subtypes. Differential expression was performed between *CD69*^+^*IGLL1*^+^ HSC-like pAML and all other pAML subtypes by the Wilcoxon rank sum test. Colors indicate significantly over-(red) or under-expressed (blue) genes by log_2_ fold change > 0.25 and adjusted p < 0.05. The x-axis represents log_2_ fold change, and the y-axis represents negative log_10_ adjusted p-value. (C) Dot plot of KEGG 2021 pathways identified as significantly enriched in *CD69*^+^*IGLL1*^+^ HSC-like pAML. The X-axis depicts the -log(adjusted p-value), the dot size depicts the number of genes in each pathway, and the dot color represents the odds ratio (higher: blue, lower: black). (D) Violin plots and UMAP feature plots of selected top differentially expressed genes. Violin plots depict normalized, scaled expression values for *CD69*^+^*IGLL1*^+^ HSC-like cells compared to all other pAML subtypes. UMAP colors indicate normalized, scaled expression (red-high, white-medium, blue-low). (E) Scatter plot of *CD69*^+^*IGLL1*^+^ gene signature score in diagnostic bone marrow biopsies from RUNX1 pAML samples in TARGET (X-axis) vs bone marrow blast percentage as measured by clinical flow cytometry (Y-axis). The correlation was assessed by the Pearson correlation test. (F) Heatmap of *CD69*^+^*IGLL1*^+^ HSC-like gene signature genes in RUNX1 samples (Y-axis) from the Beat AML bulk RNA sequencing database. Genes were selected for each subtype based on the top 30 genes by log_2_ fold change with log_2_ fold change > 0.5, adjusted p < 0.05, Wilcoxon rank sum test. Color indicates normalized, z-scaled expression value (red-high, white-medium, blue-low). Samples with signature scores (mean normalized, z-scaled for all genes) higher than the mean for all RUNX1 samples were labeled “high”, samples below the mean were labeled “low” and grouped on the Y-axis. (G) Volcano plot for *ex vivo* treatment inhibitors in the Beat AML dataset. IC_50_ values for the “high” and “low” gene signature groups were separately calculated from *ex vivo* treatment data for each treatment inhibitor using a 5-parameter log-logistic model. Then Log_2_ fold potency was calculated by log_2_(*CD69*^+^*IGLL1*^+^ high IC_50_ / *CD69*^+^*IGLL1*^+^ low IC_50_) (X-axis). Significance was tested by two-way ANOVA for the enrichment group and inhibitor concentration vs viability, p-value for the patient group effect is shown on the Y-axis. Inhibitors with absolute Log_2_ fold potency > 1.0 and p < 0.05 were considered differentially efficacious (shown in green). Inhibitors with supporting transcriptomic evidence in H-J were annotated on the plot. (H) Gene-set enrichment plot for the “WP Proteasome Degradation” pathway in *CD69*^+^*IGLL1*^+^ pAML. Bortezomib, an inhibitor with higher efficacy in *CD69*^+^*IGLL1*^+^-like samples, is a proteasome inhibitor. (I) Violin plot of *MAPK1*, a target of PD173955. (J) Gene-set enrichment plot for the “Reactome MTOR Signaling” pathway in *CD69*^+^*IGLL1*^+^ pAML. *MTOR* is a target of rapamycin, an inhibitor with less efficacy in *CD69*^+^*IGLL1*^+^-like samples. (K) Bar chart showing *CD69*^+^*IGLL1*^+^ HSC-like gene signature scores in pediatric cell lines from the GSE59808 dataset. (L) Dose-response curves for *in vitro* treatment of Kasumi-1 cells with bortezomib, rapamycin, and PD173955. UMAP: Uniform manifold approximation and projection, HSC: hematopoietic stem cell, Mono: monocyte, adjP: adjusted p-value, BM: bone marrow.

### High Risk, Treatment-resistant CD69^+^IGLL1^+^ HSC cells are predicted to be sensitive to proteasome inhibition

To identify pharmaceutical inhibitors that are specifically efficacious against cells that highly express the *CD69*^+^*IGLL1*^+^ HSC-like RUNX1 pAML gene signature, we adopted an approach similar to the Connectivity Map^35^. We first calculated enrichment for the *CD69*^+^*IGLL1*^+^ HSC-like RUNX1 pAML gene signature in RUNX1 samples from the Beat AML dataset and stratified samples into a gene signature high-expression group (score greater than average *CD69*^+^*IGLL1*^+^ score) and low-expression group (score lower than average CD69^+^IGLL1^+^ score) (**Fig 4F**). To identify inhibitors with comparatively high efficacy in *CD69*^+^*IGLL1*^+^ high samples, we compared cell viability after *ex vivo* treatment with 111 inhibitors between high and low gene-signature groups. Fifty percent inhibitory concentration (IC_50_) was estimated by a five-parameter log-logistic model and significance testing was conducted by ANOVA. Fifteen inhibitors were identified with absolute log_2_-fold potency (L_2_FP) > 1 and p-value<0.05 (**Fig 4G**). Since predictions were based on adult AML data, we further examined the pediatric scRNAseq data for transcriptomic evidence supporting predicted high or low efficacy from Beat AML *ex vivo* treatment data. Supporting evidence was considered the high expression of genes or pathways that are targeted by a pharmaceutical inhibitor predicted to be highly efficacious in Beat AML (or vice versa for low efficacy).

Bortezomib, a proteasome inhibitor, was predicted to have high efficacy (L_2_FP=2.2, p<0.05) in *ex vivo* treatment of *CD69*^+^*IGLL1*^+^ high samples compared to *CD69*^+^*IGLL1*^+^ low samples (**Fig 4G**). This is further supported by the enrichment of the proteasome degradation pathway in the *CD69*^+^*IGLL1*^+^ pAML single cells (**Fig 4H**). Thus, the higher efficacy predicted in Beat AML samples with high signature scores for the *CD69*^+^*IGLL1*^+^ HSC-like pAML subtype was supported by enrichment for the corresponding pathway in scRNAseq. By similar logic, rapamycin (L_2_FP=-7.60, p<0.05) and PD173955 (L_2_FP=-2.48, p<0.05) had negative L_2_FP, indicating minimal impact on the viability of cells expressing the *CD69*^+^*IGLL1*^+^ HSC-like pAML signature (**Fig 4G**). *MAPK1*, a target of PD173955, had no expression in 70.8% of *CD69*^+^*IGLL1*^+^ HSC-like pAML cells (**Fig 4I**). Similarly, the MTOR (mechanistic target of rapamycin) signaling pathway was not significantly enriched in *CD69*^+^*IGLL1*^+^ HSC-like pAML (p=0.49, **Fig 4J**). Therefore, the predicted lower efficacy in samples with high *CD69*^+^*IGLL1*^+^ signature scores was supported by a lack of expression or enrichment for the targets of PD173955 and rapamycin.

To further validate these findings, we conducted *in vitro* validation experiments using the Kasumi-1 cell line. Kasumi-1 is derived from a RUNX1-RUNX1T1 patient and expresses the highest mean z-score for the *CD69*^+^*IGLL1*^+^ gene signature from an array of pAML cell lines (**Fig 4K**)^36^. *In vitro* treatment of Kasumi-1 with bortezomib, the predicted high-efficacy inhibitor, depicted an IC_50_ of 3.59 nM with a 95% confidence interval (CI) of 3.42-3.79 (**Fig 4L**). The predicted lower efficacy inhibitors, PD173955 (IC_50_=56.2 nM, 95% CI: 25.4-87.0), and rapamycin (did not reach IC_50_ up to 118 nM) were considerably less efficacious than bortezomib (**Fig 4L**). These results indicate *CD69*^+^*IGLL1*^+^ pAML is enriched for proteasome degradation and sensitive to bortezomib, presenting a potentially targeted chemotherapeutic for HRTR RUNX1 pAML.

***High-risk, treatment-resistant CD69^+^IGLL1^+^ HSC cells promote an inflammatory, exhausted immune microenvironment*.**

Given the implicated immune interactions in *CD69*^+^*IGLL1*^+^ HSC-like pAML gene expression, our next objective was to characterize the lymphoid cell changes in samples enriched for this pAML subtype. As *CD69*^+^*IGLL1*^+^ HSC-like pAML was a RUNX1 subtype, we first merged, batch corrected, and clustered the lymphoid cells from RUNX1 samples (**Fig 5A**). Lymphoid cell types were annotated using canonical markers for major subtypes (e.g. naïve, memory, cytotoxic) including CD4^+^ and CD8^+^ T cells, NK, and B cells (**Fig 5B**). To estimate the impact of *CD69*^+^*IGLL1*^+^ HSC-like pAML on lymphoid cells, samples were stratified into “*CD69*^+^*IGLL1*^+^” if *CD69*^+^*IGLL1*^+^ cells had the highest count among malignant cells, or “other” for any other pAML subtype (**Fig 5C**). In *CD69*^+^*IGLL1*^+^ predominant samples, CD8^+^ T cells expressed markers of exhaustion at higher levels as compared to other samples (*LAG3* Log_2_FC=0.22, adjusted p<0.05, *BATF* Log_2_FC=0.20, adjusted p<0.05, **Fig 5D**). Comparison of the proportion of lymphoid cell types in *CD69*^+^*IGLL1*^+^ versus other samples showed CD8^+^ cytotoxic T cells had significantly higher abundance (p<0.05), while CD4^+^ memory T cells had significantly lower abundance within *CD69*^+^*IGLL1*^+^ samples (p<0.05, **Fig 5E**). To validate if the same trends were manifested in RUNX1 pAML TARGET samples, deconvolution analysis was performed. Consistent with scRNAseq, CD8^+^ T cytotoxic gene scores were positively correlated (R=0.22, p<0.05) and CD4^+^ memory gene scores were negatively correlated with *CD69*^+^*IGLL1*^+^ scores (R=-0.15, p<0.05, **Fig 5F-G**). These results indicate the association of *CD69*^+^*IGLL1*^+^ HSC-like pAML with exhaustion in CD8^+^ T cells as well as the accumulation of CD8^+^ cytotoxic T-effector cells and depletion of CD4^+^ memory T cells.

**Figure 5:**
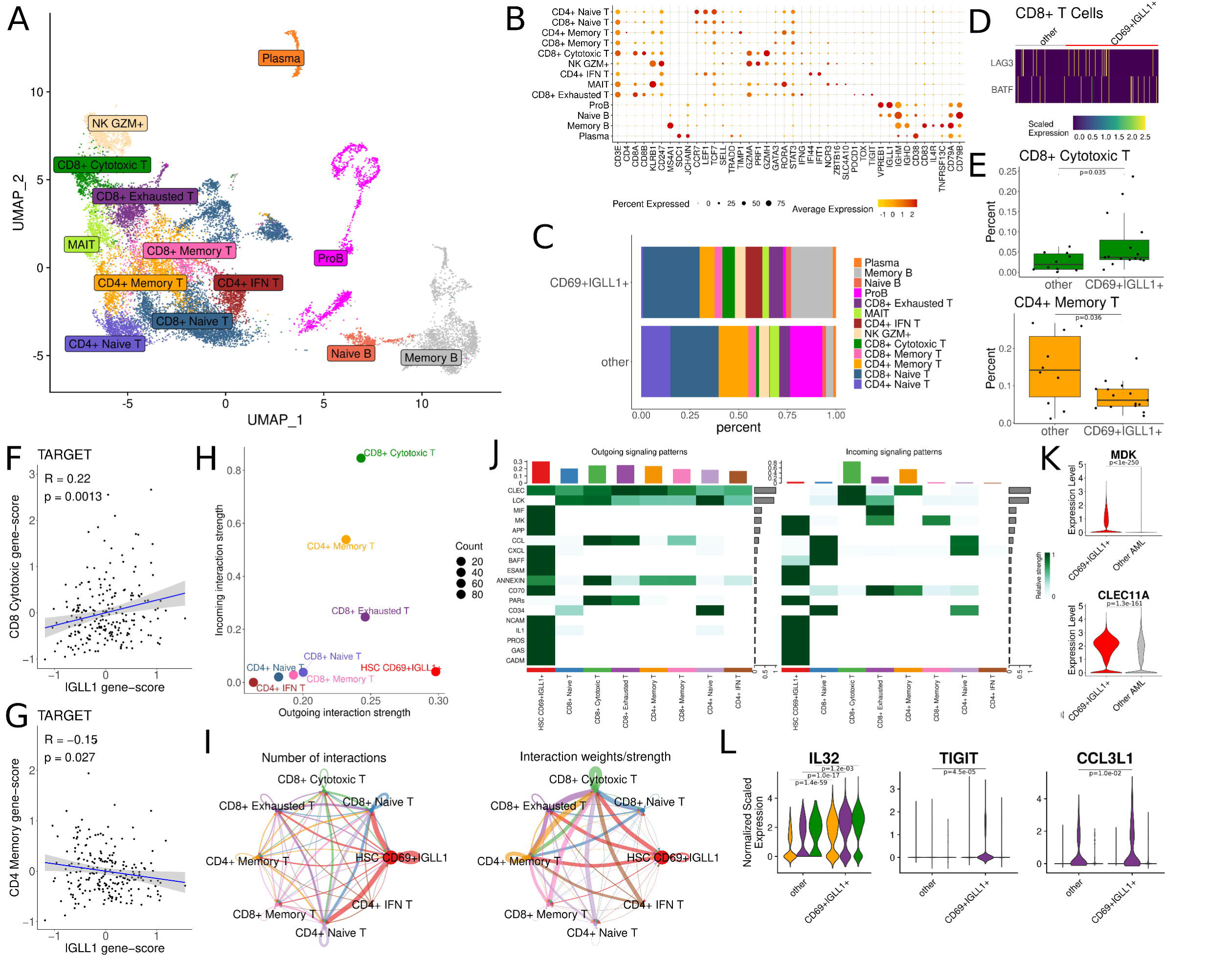
High-risk, treatment-resistant hematopoietic stem cell (HSC)-like cells in RUNX1 patients promote an inflammatory, exhausted immune microenvironment. (A) UMAP embeddings for lymphoid cells from RUNX1 pAML samples. Cells are colored by lymphoid subtype annotation. Annotations were assigned based on canonical marker gene expression (B) Dot plot showing expression of canonical lymphoid subset marker genes. Markers used included: Naïve T (*CCR7*, *TCF7, LEF1*), memory (*SELL*, *TRADD*, *TIMP*), cytotoxic (*GZMA*, *PRF1*, *GZMH*), interferon (IFN) stimulated (*IFI44*, *IFIT1*), mucosal-associated invariant T cell (MAIT) (*NCR3*, *ZBTB16*, *SLC4A10*), exhausted (*PDCD1*, *TOX*, *TIGIT*), pro-B (*VPREB1*, *IGLL1*), naïve B (increased *CD79B*/*CD79A* ratio, increased *IGHM*/other IGH ratio), memory B (*CD83*, *IL4R*, *TNFRSF13C*, decreased *CD79B*/*CD79A* ratio), and plasma (*CD38*). The color intensity represents gene expression (red-high, yellow-low), and the dot size represents the percentage of cells expressing each gene within individual cell types. (C) Bar plot of the relative proportion of lymphoid subtypes between samples in which *CD69*^+^*IGLL1*^+^ HSC-like pAML was the malignant subtype with the highest cell count and samples with any other subtype as the most abundant malignant subtype. (D) Heatmap of exhaustion markers, *BATF* and *LAG3*, in CD8^+^ T cells. Cells are grouped along the Y-axis by *CD69*^+^*IGLL1*^+^ samples vs other samples. Both markers were expressed at higher levels in *CD69*^+^*IGL11*^+^ samples (Wilcoxon rank sum test, adjusted p < 0.05). Relative gene expression is indicated by color with yellow as highly expressed and purple as lowly expressed. (E) Box plots for the proportion of lymphoid cells annotated as CD8^+^ cytotoxic (top) or CD4^+^ memory (bottom) T cells compared between *CD69*^+^*IGLL1*^+^ and other samples. Significance testing performed by student’s t-test. (F-G) Scatter plot of *CD69*^+^*IGLL1*^+^ gene signature score in RUNX1 bulk RNA sequencing samples in TARGET versus canonical gene signature scores of CD8^+^ cytotoxic T cells (F) and CD4^+^ memory T cells (G). Gene signature scores were calculated by the averaged, normalized, z-scaled expression of all genes within respective gene scores. The correlation coefficient and significance between *CD69*^+^*IGLL1*^+^ scores and the respective T-cell scores were calculated using Pearson correlation. (H) Scatter plot of the Outgoing Interaction Strength and Incoming Interaction Strength metrics produced by CellChat colored and labeled by cell type. Cell types to the far right of the graph are considered to have strong outgoing interactions by CellChat metrics and cell types towards the top are considered to have strong incoming interactions. (I) Circos plots indicating the relative number and strength of communications between cell types identified by CellChat. Chord color indicates the cell type sending signals and width indicates the number (left) and strength (right) of communication between connected cell types based on ligand and receptor correlation. (J) Heatmap of the pathways with the highest interaction weights in *CD69*^+^*IGLL1*^+^ pAML and T cells. Rows correspond to sending cell types, columns correspond to receiving cell types. Darker colors indicate stronger interactions. (K) Violin plots of midkine (*MDK*) and C-lecithin 11A (*CLEC11A*), genes associated with the top two pathways by outgoing interaction strength in *CD69*^+^*IGLL1*^+^ HSC-like cells by CellChat. Comparison between *CD69*^+^*IGLL1*^+^ HSC-like pAML and all other pAML subtypes was executed by the Wilcoxon rank sum test. (L) Violin plots of *IL32*, a marker of chronic inflammation in cancer, and two markers of exhaustion (*TIGIT*, *CCL3L1*). Violins are colored by the T cell subtype and compared between *CD69*^+^*IGLL1*^+^ and other samples by the Wilcoxon rank sum test. UMAP: Uniform manifold approximation and projection, HSC: hematopoietic stem cell.

To further study the interaction of *CD69*^+^*IGLL1*^+^ HSC-like pAML with T cells in RUNX1 samples we performed cellular communication analysis using ligand and receptor expression by CellChat^37^. The analysis showed that *CD69*^+^*IGLL1*^+^ HSC-like pAML cells exhibit the strongest outgoing interactions, with CD8^+^ cytotoxic, CD4^+^ memory, and CD8^+^ exhausted T cells displaying the strongest incoming signals (**Fig 5H-I**). Midkine (MK/MDK), a growth factor associated with treatment resistance in AML^38^ and T cell dysfunction in other cancers^39,40^, emerged as one of the strongest outgoing signals in *CD69*^+^*IGLL1*^+^ pAML and among the strongest incoming signals in CD8^+^ exhausted T cells (**Fig 5J**). *MDK* was overexpressed in *CD69*^+^*IGLL1*^+^ pAML relative to other subtypes (Log_2_FC: 0.95, adjusted p<0.05, **Fig 5K**). Additionally, C-lecithins (CLEC), a family of growth factor glycoproteins known to associate with residual disease in AML^9,41^, were also identified as one of the top outgoing pathways in *CD69*^+^*IGLL1*^+^ pAML, as the top incoming pathway to CD8^+^ cytotoxic and CD4^+^ memory T cells, and expressed at higher levels in *CD69*^+^*IGLL1*^+^ pAML versus other subtypes (Log_2_FC: 0.29, adjusted p<0.05, **Fig 5J-K**).

To further explore differences in expression in T cell subsets from in *CD69*^+^*IGLL1*^+^ samples we performed DE of CD8^+^ cytotoxic, CD4^+^ memory, and CD8 exhausted T cells between *CD69*^+^*IGLL1*^+^ and other RUNX1 samples. All three T subsets had increased *IL32*, a marker of chronic inflammation^42^ (L_2_FC of 1.39, 0.70, 0.99 respectively, all adjusted p<0.05). CD8^+^ exhausted T cells also had increased markers of exhaustion (*TIGIT*: L_2_FC=0.56 and *CCL3L1*: L_2_FC=1.16, both adjusted p<0.05, **Fig 5K**)^43^.

In summary, *CD69*^+^*IGLL1*^+^ HSC-like pAML appears to promote chronic inflammation and exhaustion of T cells through the CLEC and MDK pathways, underscoring their potential role in disease progression and immune evasion.

### High-risk, treatment-resistant HSC-like cells in FLT3 patients express leukemic stem cell markers and utilize antioxidant metabolism

To study how HRTR pAML in FLT3 patients promotes poor outcomes, we generated the transcriptome profiles (absolute L_2_FC > 0.25, adjusted p<0.05) of *CD69*^+^*TRGC2*^+^ HSC-like pAML with 74 significantly over-expressed and 152 downregulated genes (**Fig 6A-B, Supplemental Table 7**). *TRGC2*, an isotype of the T cell receptor alternate reading frame protein (TARP) previously identified as a marker of LSCs in FLT3 adult AML^44^, was highly expressed (L_2_FC=1.18, adjusted p<0.05) as well as immature markers including *PRSS57* (L_2_FC=0.87, adjusted p<0.05). Additionally, malignant cells overexpressed genes from the pediatric Leukemic Stem Cell set,^11^ including *SPINK2* (L_2_FC=1.98, adjusted p<0.05) and *FAM30A* (L_2_FC=1.12, adjusted p<0.05, **Fig 6B**), both associated with poor outcomes. Pathway analysis identified the enrichment of drug metabolism as well as glutathione metabolism, a mediator of reactive oxygen species processing, in *CD69*^+^*TRGC2*^+^ pAML (**Fig 6C**)^45^. Glutathione metabolism has previously been described as a resistance mechanism to anthracyclines, one of the two first-line pAML chemotherapies^46,47^. Two glutathione pathway genes known to protect cells from oxidative stress, *MGST1* (L_2_FC=0.74, adjusted p<0.05) and *MGST3* (L_2_FC=0.26, adjusted p<0.05) were highly expressed (**Fig 6D**)^45^. Thus, *CD69*^+^*TRGC2*^+^ HSC-like pAML is enriched for known LSC genes and appears to utilize antioxidant metabolism to support persistence through treatment.

**Figure 6:**
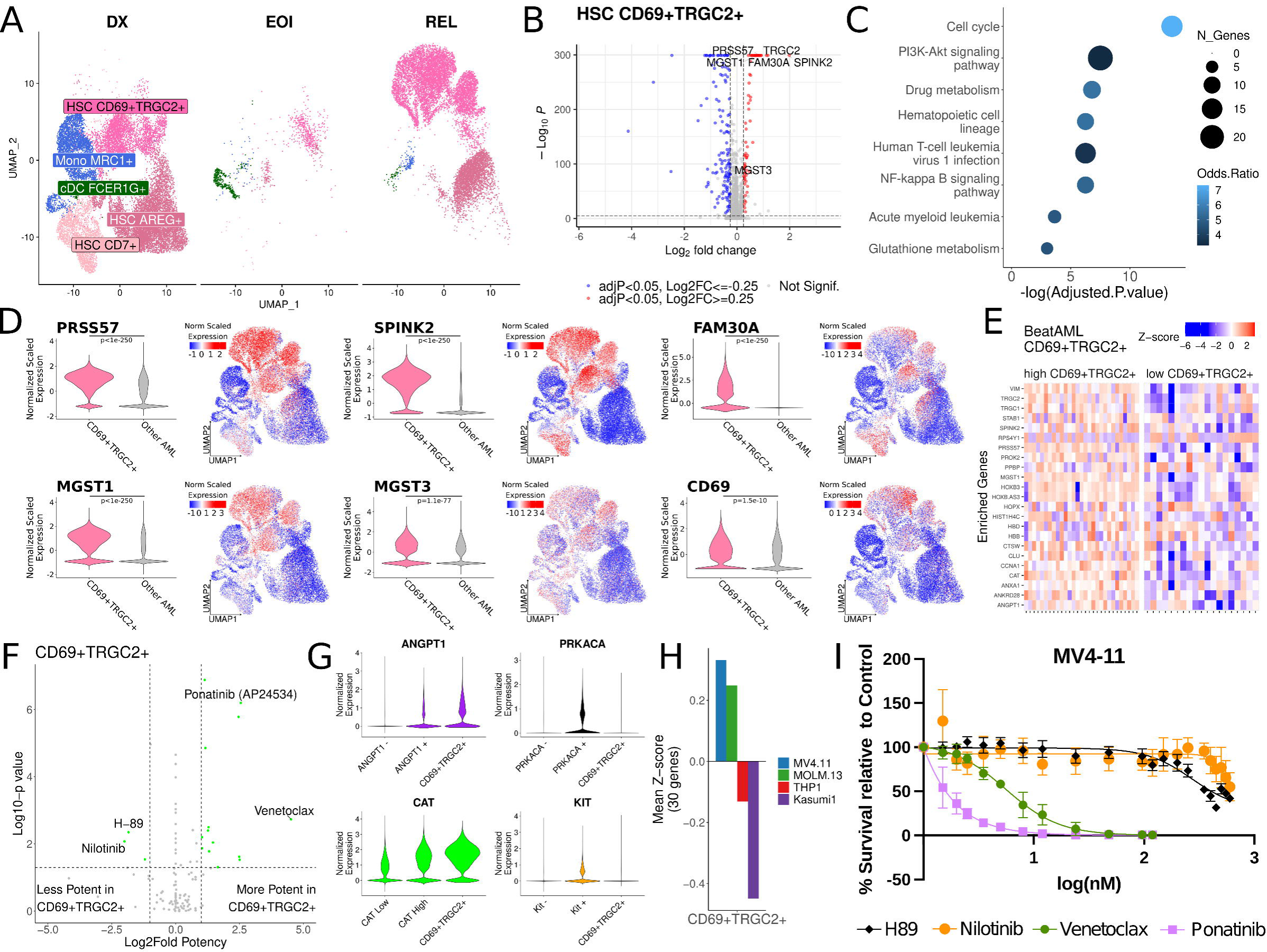
High-risk, treatment-resistant HSC-like cells in FLT3 patients express leukemic stem cell markers and utilize antioxidant metabolism. (A) UMAP embeddings depicting the FLT3 pAML cells at diagnosis (DX) end-of-induction (EOI) and relapse (REL). (B) Volcano plot of differentially expressed genes in *CD69*^+^*TRGC2*^+^ HSC-like pAML vs all other subtypes. Differential expression was performed between *CD69*^+^*TRGC2*^+^ HSC-like pAML and all other pAML subtypes by the Wilcoxon rank sum test. Colors indicate significantly over-(red) or under-expressed (blue) genes by log_2_ fold change > 0.25 and adjusted p < 0.05. The x-axis represents log_2_ fold change, and the y-axis represents negative log_10_ adjusted p-value. (C) Dot plot of GO Biological pathways identified as significantly enriched in *CD69*^+^*TRGC2*^+^ HSC-like pAML. The X-axis depicts the -log(adjusted p-value), the dot size depicts the number of genes in each pathway, and the dot color represents the odds ratio (higher: blue, lower: black). (D) Violin plots and UMAP feature plots of selected top differentially expressed genes in *CD69*^+^*TRGC2*^+^ cells. Violin plots depict normalized, scaled expression values for *CD69*^+^*TRGC2*^+^ HSC-like cells compared to all other pAML subtypes. UMAP colors indicate normalized, scaled expression (red-high, white-medium, blue-low). (E) Heatmap of *CD69*^+^*TRGC2*^+^ HSC-like gene signature genes in FLT3 samples (Y-axis) from the Beat AML bulk RNA sequencing database. Genes were selected based on the top 30 genes by log_2_ fold change with log_2_ fold change > 0.5, adjust p < 0.05, Wilcoxon rank sum test. Color indicates normalized, z-scaled expression value (red-high, white-medium, blue-low). Samples with signature scores (mean normalized, z-scaled for all genes) higher than the mean for all FLT3 samples were labeled “high”, samples below the mean were labeled “low” and grouped on the X-axis. (F) Volcano plot for *ex vivo* treatment inhibitors in the Beat AML dataset. IC_50_ values for the “high” and “low” gene signature groups were separately calculated from *ex vivo* treatment data for each treatment inhibitor using a 5-parameter logistic model. Then Log_2_ fold potency was calculated by log_2_(*CD69*^+^*TRGC2*^+^ high IC_50_ / *CD69*^+^*TRGC2*^+^ low IC_50_) (x axis). Significance was tested by two-way ANOVA for the enrichment group and inhibitor concentration vs viability, the p-value for the patient group effect is shown on the Y-axis. Inhibitors with supporting transcriptomic evidence in G were annotated on the plot. (G) Violin plot of *ANGPT1*, a target of ponatinib, *CAT*, a gene that predicts sensitivity to venetoclax, *PRKACA*, a target of H-89, and *KIT*, a target of nilotinib. (H) Bar chart showing *CD69*^+^*TRGC2*^+^ HSC-like gene signature scores in pediatric cell lines from the GSE59808 dataset. (I) Dose-response curves for *in vitro* treatment of Kasumi-1 cells with ponatinib, venetoclax, nilotinib, and H-89. HSC: hematopoietic stem cell, Mono: monocyte, cDC: classical dendritic cell, adjP: adjusted p-value.

### High-risk, treatment-resistant FLT3 HSC-like cells are sensitive to BCL2 and kinase inhibition

To predict pharmaceutical inhibitors for targeting malignant cells expressing the *CD69*^+^*TRGC2*^+^ HSC-like gene signature, we analyzed the BeatAML dataset as described for RUNX1 pAML. We deconvoluted Beat AML’s bulk RNA sequencing of FLT3 mutated samples to stratify samples into “high” and “low” *CD69*^+^*TRGC2*^+^ gene signature groups (**Fig 6E**). Comparison of cell viability in *ex vivo* treatment data between samples labeled high and low identified seventeen inhibitors with significantly higher or lower efficacy in samples enriched for the *CD69*^+^*TRGC2*^+^ HSC-like pAML signature (absolute L_2_FP > 1.0 and p<0.05, **Fig 6E**). Ponatinib, a multi-kinase inhibitor that targets *ANGPT1*, was highly efficacious in samples enriched for the *CD69*^+^*TRGC2*^+^ gene signature (L_2_FP=2.550, p<0.05, **Fig 6F**). The relatively high efficacy of ponatinib was supported by the high expression of *ANGPT1* in *CD69*^+^*TRGC2*^+^ pAML cells (L_2_FC=0.83, p<0.05, **Fig 6G**). Venetoclax was also more efficacious in high *CD69*^+^*TRGC2*^+^ samples (L_2_FP=4.52, p<0.05) and supported by overexpression of *CAT*, a gene associated with venetoclax sensitivity^48^, (L_2_FC=0.85, p<0.05, **Fig 6G**). H-89 (L_2_FP=-1.84) and nilotinib (L_2_FP=-2.00) were predicted to be less efficacious in samples with high *CD69*^+^*TRGC2*^+^ scores. The low efficacy was supported by the lack of expression of their targets: *PRKACA* (0 counts in 92.7% of cells) and *KIT* (0 counts in 82.1% of cells) respectively (**Fig 6G**). *TRGC1* and *TRGC2* were previously found to be highly expressed in MV4-11 cells, which harbor FLT3-ITD mutation^44^. MV4-11 scored the highest similarity to *CD69*^+^*TRGC2*^+^ gene-signature (**Fig 6H**)^49^. Consistent with the Beat AML findings, *in vitro* treatment of MV4-11 found ponatinib (IC_50_: 0.19 nM, 95% CI: 0.00-1.28) and venetoclax (IC_50_=5.56 nM, 95% CI: 4.59-6.51) highly efficacious (**Fig 6I**). Conversely, H-89 (IC_50_: 272.7 nM, 95% CI: 208.5-6710.0) and nilotinib were found to be minimally efficacious (IC50: 565.3 nM, 369.0-10000.0, **Fig 6I**). Thus, *CD69*^+^*TRGC2*^+^ pAML cells express a gene signature associated with increased *ex vivo* and *in vitro* sensitivity to venetoclax and ponatinib. Additionally, these cells show high expression levels of known targets and sensitivity genes for these inhibitors, indicating that venetoclax and ponatinib are particularly effective for FLT3 HRTR pAML.

### Core binding factor mutated pAML patients depict elevated mast cells associated with poor outcomes

Mast cells, tryptase-producing mature myeloid cells, typically make up <1% of the BM microenvironment and are not well characterized in pAML^13^. Our subtype annotations (**Fig 3D**) identified pAML cells with > 4.0 log_2_-fold expression of tryptase genes *TPSAB1*, *TPSB2*, and mast markers *IL5RA* (L_2_FC=2.53) and *FCER1A* (L_2_FC=2.13, all adjusted p<0.05, **Fig 7A-B**, **Supplemental Table 8**). Consistent with adult AML studies^14^, mast cells were almost exclusively identified in RUNX1 or CBFB samples, collectively the CBF pAMLs. Approximately 29% of CBF samples (n=24) had >5% mast cells versus 1.7% (n=58) of FLT3, KMT2A, and “other” samples (**Fig 7C**). To validate if mast cells were CBF-enriched in TARGET pAML data also, we calculated enrichment for mast cells using a canonical gene signature (*TPSAB1*, *TPSB2*, *IL5RA*, *FCER1A*) in diagnostic BM samples. CBF pAML had significantly higher enrichment than pAML from other cytogenetic backgrounds (p<0.05, **Fig 7D**). Together, our scRNAseq and TARGET cohorts demonstrate mast cells are preferentially enriched in CBF pAML.

**Figure 7:**
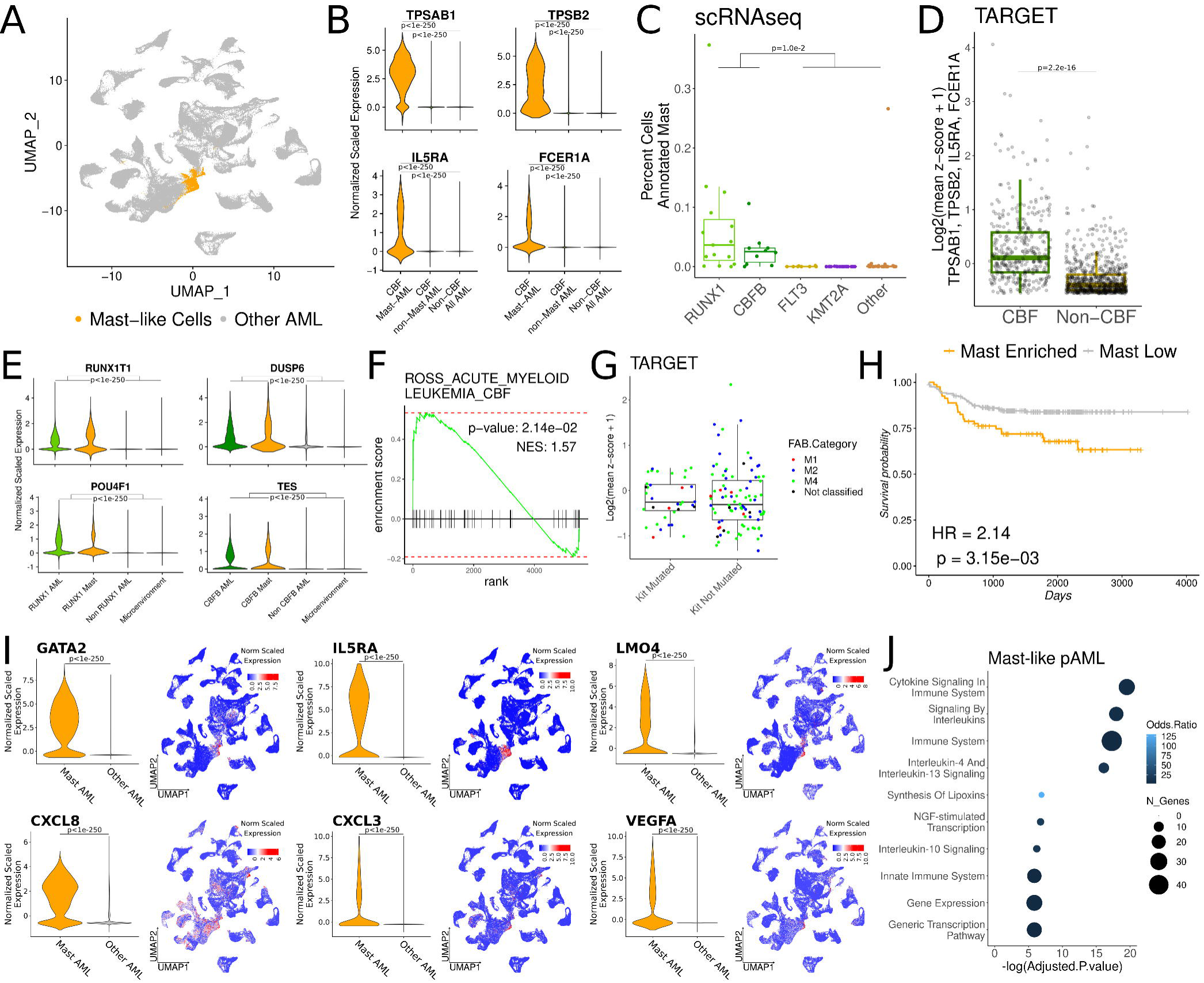
Core binding factor (CBF) mutated pAML patients depict elevated mast cells associated with poor outcomes. (A) UMAP embeddings of malignant pAML cells from matched samples. Colors highlight the mast cells (orange) versus all other pAML (gray). (B) Violin plots for canonical mast cell marker genes. Normalized expression was compared between mast cells in CBF samples, non-mast malignant cells in CBF samples, and malignant cells from all other cytogenetic backgrounds using the Wilcoxon rank sum test with a significance threshold of 0.05. (C) Box plots of the proportion cells annotated as mast cells in each cytogenetic group. Each point represents the proportion of cells in a single patient. Mast proportions were compared between the CBF patients (CBFB and RUNX1) and all non-CBF cytogenetic groups (FLT3, KMT2A, other) by t-test. (D) Box plot of mast enrichment scores for CBF and non-CBF bulk RNA sequencing samples from the TARGET database. Mast enrichment scores were calculated by Log_2_(mean z-score + 1) for *TPSAB1*, *TBSP2*, *FCER1A*, and *IL5RA* and compared by student’s t-test with a significance threshold of 0.05. (E) Violin plot of genes known to be markers of RUNX1 pAML (*RUNX1T1*, *POU4F1*) and CBFB pAML (*DUSP6*, *TES*). Expression is compared between non-mast pAML from either RUNX1 or CBFB, mast cells, pAML from other cytogenetic groups, and non-malignant cells (e.g. T and B cells). Statistical significance was tested by Wilcoxon rank sum. (F) Gene set enrichment for the Ross et al. CBF AML gene list based on the ranked ordered gene list of mast cells in CBF patients vs non-malignant cells from CBF patients. Enrichment indicates that, compared to non-malignant cells, mast cells express a CBF pAML-like gene signature. (G) Box plot of mast enrichment values in CBF bulk RNAseq samples in TARGET. Kit mutation is a known driver of systemic mastocytosis, a different myeloid neoplasm of mast cells. The mast enrichment score in CBF samples was compared between Kit mutated and Kit not mutated samples by student’s t-test. The comparable mast scores between Kit mutated and Kit not mutated samples indicate that the relative mastocytosis in CBF patients does not appear to be confounded by Kit mutation. (H) Kaplan Meier curve comparing overall survival between CBF TARGET samples labeled as mast enriched or mast low. Samples were considered mast enriched for a gene signature score two standard errors above the mean. Twenty-seven percent of TARGET samples were labeled mast enriched, comparable to the 29% of samples found to be mast enriched in scRNAseq. Overall survival was compared by the Cox proportional hazards model. (I) Violin plots and UMAP feature plots of selected top differentially expressed genes. Violin plots depict normalized, scaled expression values for mast pAML cells compared to all other pAML subtypes. UMAP colors indicate normalized, scaled expression (red-high, white-medium, blue-low). (J) Dot plot of Reactome pathways identified as significantly enriched in mast-like pAML. The X-axis depicts the -log(adjusted p-value), the dot size depicts the number of genes in each pathway, and the dot color represents the odds ratio (higher: blue, lower: black). scRNAseq: Single-cell RNA sequencing, FAB: French-American-British classification, UMAP: uniform manifold approximation for projection.

To further evaluate if gene expression of mast cells depicts any similarity with malignant pAML cells, we adopted two complementary approaches. First, we examined the expression of established markers of RUNX1 (*RUNX1T1*, *POU4F1*^50,51^) and CBFB (*DUPS6*, *TES*^51^) pAML. Mast cells expressed all four genes at comparable levels to pAML from RUNX1/CBFB patients respectively, and higher than pAML from other cytogenetic backgrounds and microenvironment cells (**Fig 7E**). Second, we tested mast cells for the enrichment of the CBF pAML gene set published by Ross et al^51^. Gene set enrichment analysis on mast cells showed significant enrichment for the ROSS_ACUTE_MYELOID_LEUKEMIA_CBF gene set (NES: 1.57, p<0.05, **Fig 7F**) as compared to other non-malignant cells in CBF pAML samples. This supports the insight that mast cells in CBF pAML are malignant. Systemic Mastocytosis (SM) is a neoplasm of mast cells driven by KIT mutation in >90% of cases^52^ that can co-occur with pAML. In our scRNAseq patient cohort, 1/24 CBF patients had documented *KIT* mutation (**Supplemental Table 2**). To determine if *KIT* mutation influenced mast abundance in TARGET pAML data, we compared mast cell enrichment scores between *KIT*-mutated and non-mutated samples and observed no significant differences (p=0.92, **Fig 7G**). Thus, it appears that mast cells in CBF pAML are components of the tumor ecosystem independent of *KIT* mutation.

To determine if malignant mast cell enrichment was associated with worse outcomes in TARGET pAML data, we stratified patients based on malignant mast cell gene signature enrichment score (*TPSAB1*, *TPSB2*, *IL5RA*, *FCER1A*, and *CD69* which was highly expressed in both mast-like pAML subtypes). Two standard errors above the mean stratified 26.0% (99/381) of CBF samples in TARGET as “mast-high” (versus 29.2% of samples in scRNAseq). Mast cell enrichment was associated with shortened overall survival (HR=2.14, p<0.05, **Fig 7H**). To further characterize malignant mast cells, we generated marker genes by comparing the expression profile of mast cells vs all other cell malignant pAML cell subtypes. Mast cells overexpressed genes associated with immaturity and poor survival: *GATA2* (Log_2_FC=2.63, p<0.05)^53,54^, *IL5RA* (Log_2_FC=2.58, p<0.05)^55,56^, and *LMO4* (Log_2_FC=2.16 p<0.05)^55^ (**Fig 7I**). Mast cells also overexpressed genes associated with tumor survival and immune signaling, including *CXCL8* (Log_2_FC=2.53, p<0.05) and *CXCL3* (Log_2_FC=1.32, p<0.05)^57^, as well as the pro-angiogenic factor *VEGFA* (Log_2_FC=1.37, p<0.05)^58,59^ (**Fig 7I**). Consistent with the high expression of immune mediating genes, enriched Reactome pathways included cytokine signaling, immune system function, and IL-4 and IL-13 signaling (**Fig 7J**). Thus, mast cells express genes consistent with a mixed mature and immature phenotype and immune microenvironment mediating factors. In summary, single-cell analysis provided novel insight into the association of mast cells with CBF pAML and poor outcomes in a patient cohort typically associated with standard risk.

## Discussion

The innovation of single-cell technologies enabled the interrogation of the heterogeneity of pAML cells and uncovered new insights into treatment resistance and interaction with the immune microenvironment^6–9^. Herein we report the largest combined scRNAseq analysis of pAML to date, including 708,285 cells from 164 BM biopsies obtained from 95 patients. The large cohort enabled the dissection of malignant cell heterogeneity across cytogenetic categories and paired analysis among samples collected at the time of disease diagnosis, end of induction, and relapse. Critically, this enabled the identification of HRTR pAML subtypes that were unique to RUNX1, FLT3, and CBFB pAML. We observed that both immature and well-differentiated pAML subtypes were associated with treatment resistance and poor outcome. Excitingly, this is the first report of mast-like malignant cells in CBF pAML. Thus, it appears that both immature and mature HRTR pAML cell subtypes contribute to poor outcomes for pAML patients.

Existing risk stratification algorithms use cytogenetic mutations to inform pre-treatment risk. However, as many as 30% of standard-risk patients relapse within 5 years, and up to 50% of high-risk patients sustain remission beyond 5 years^60^. Thus, there is a need to identify biomarkers to differentiate patients within existing cytogenetic categories. Leukemic stem cells are considered a key driver of relapse in pAML^10,11^. Previous studies have identified potential LSC markers in cohorts of pAML from mixed cytogenetic backgrounds including *CD69*, *CLEC12A*, and gene sets such as the LSC6 and LSC17^11,12,61,62^. Our findings build on prior work by uncovering LSC markers from HRTR subtypes specific to RUNX1 and FLT3 pAML. RUNX1 HRTR overexpress *CD69* and *IGLL1*, a gene typically associated with B cells, that has been reported as enriched in relapsed pAML^8^. FLT3 HRTR pAML overexpressed *CD69* and *TRGC2*, an isotype of the TARP protein previously described as a poor prognostic indicator and LSC marker in FLT3 pAML^44,63^. *CD69*^+^*IGLL1*^+^ HSC-like pAML manifests resistance by over-expressing genes related to proliferative and immune remodeling pathways, while *CD69*^+^*TRGC2*^+^ HSC-like pAML utilizes antioxidant metabolism. Together with our findings, there is compelling evidence that *CD69*, *IGLL1* (RUNX1), and *TRGC2* (FLT3) could serve as biomarkers of high-risk disease. Future studies will be needed to evaluate if prospective identification of HRTR pAML cells with these surface markers is associated with early relapse and persistence through chemotherapy.

Analysis of the Beat AML adult *ex vivo* treatment data predicted inhibitors specifically efficacious in patients with AML enriched for the gene signatures of *CD69*^+^*IGLL1*^+^ and *CD69*^+^*TRGC2*^+^ HSC-like pAML. We provided pediatric evidence for selected inhibitors by finding gene expression in scRNAseq that supported the predicted efficacy, with subsequent validation *in vitro* using pediatric cell lines. Three inhibitors were found to be potentially specifically efficacious in HRTR pAML. Ponatinib, a multi-kinase inhibitor, has not been studied in pAML but has been studied in Philadelphia chromosome (BCR-ABL, one of ponatinib’s targets) positive pediatric leukemia^64,65^. Venetoclax has ongoing clinical studies in pAML, with some promising results in relapsed/refractory pAML^66,67^. Our findings suggest that both may have specific uses in targeting HRTR FLT3 pAML. Interestingly, the AML1031 study found bortezomib with standard chemotherapy for *de novo* pAML produced no difference in overall survival in a cohort of mixed cytogenetics^68^. Our findings indicate that bortezomib may be more efficacious in RUNX1 pAML with *CD69*^+^*IGLL1*^+^ HSC-like cells as compared to pAML agnostic to the cytogenetic background. Further studies are needed to characterize HRTR pAML from BM biopsies for *ex vivo* treatment with bortezomib, ponatinib, and venetoclax to determine if comparable efficacy is observed in patient-derived pediatric samples. This study provides an exciting foundation for the prospective identification of HRTR pAML and precision treatment to improve their clearance from the bone marrow.

This is the first large molecular study examining mast cells in pAML. Mast cells were found almost exclusively in CBF pAML patients, persisted through chemotherapy, and were associated with relapse and worse overall survival despite CBF mutations generally classifying as standard risk. Mast cell persistence through chemotherapy is consistent with case reports of systemic mastocytosis with an associated myeloid neoplasm (SM-AMN), in which patients who also had CBF pAML were found to have persistent mast cells after induction^16,19,69–71^. Intriguingly, SM-AMN patients with CBF mutations survived less than two years in most cases, though larger studies are needed to validate if such poor outcomes are validated in a larger cohort and CBF pAML patients without SM. Studies of adult CBF AML also reported elevated mast cells, persistence through chemotherapy, and sustained elevations in tryptase predicted early relapse for patients in apparent remission^14,15,72^. It appears that mast-like pAML acts as a microenvironmental mediator, expressing high levels of tryptase, cytokines, and pro-angiogenic factors. Further studies are needed to define how malignant mast cells contribute to pAML chemoresistance and design treatments to enhance their clearance.

This study demonstrates that both immature and mature cell types contribute to relapse in pAML. The unique gene expression of HRTR cells provided foundational insight into deranged biological functions and identified potential targets for prospective identification and treatment. Determining the specificity of these cell surface genes for HRTR pAML subtypes at the protein level, as well as *ex vivo* drug sensitivity testing, are critical next steps. Importantly, this is the first study to associate malignant mast cells with poor outcomes in CBF pAML. Targeting mast cells has the potential to enhance the durability of remission; future studies will explore how these cells promote pAML chemoresistance. Further exploration using spatial transcriptomics of BM biopsies from CBF pAML patients will shed light on the spatial localization and communication landscape of mast cells in the tumor microenvironment. Together these findings provide an exciting foundation for ongoing work examining the contribution of immature and mature myeloid cells to relapse and how these cells can be treated to improve patient outcomes.

---

**Figure S1:**
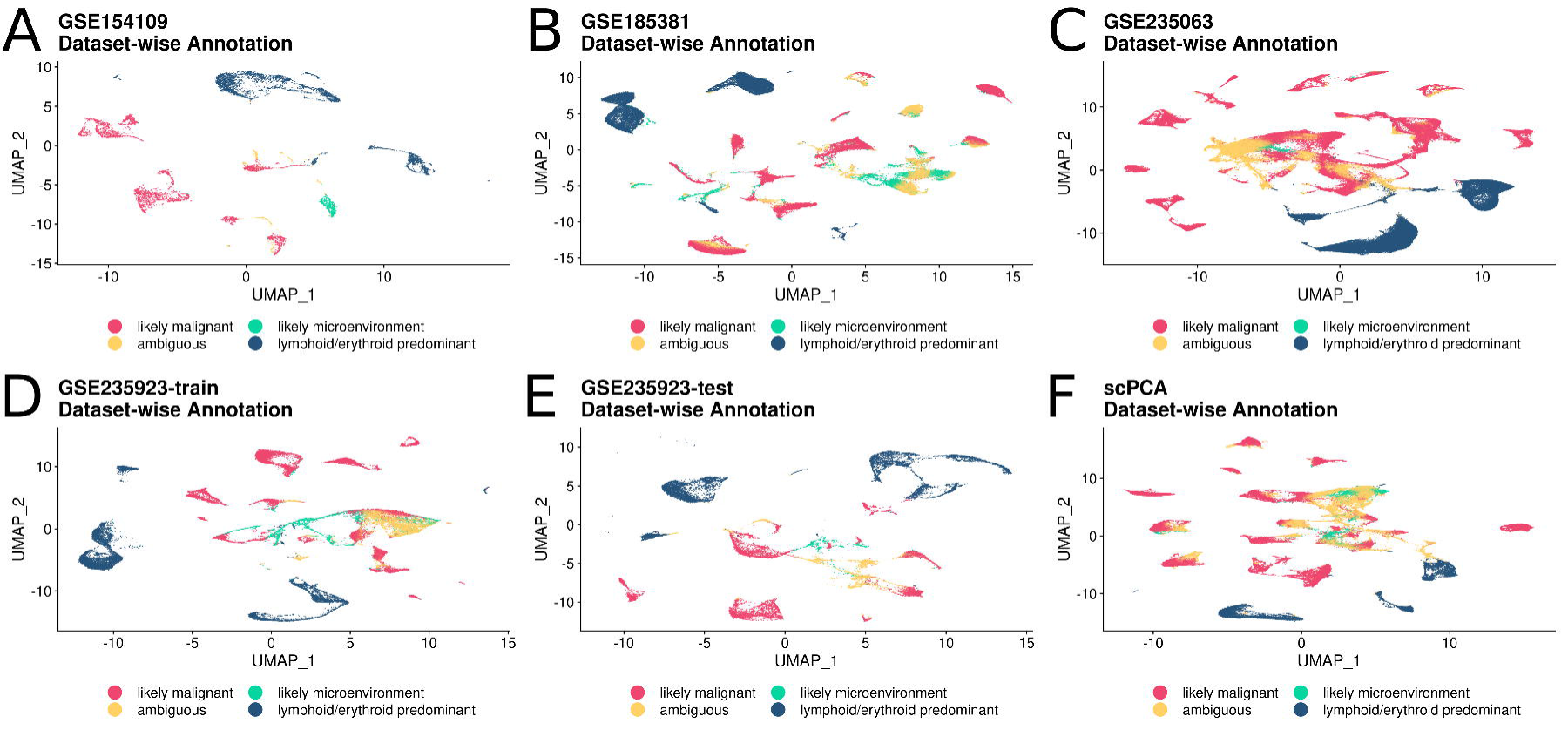
Initial dataset-wise annotation of malignant and microenvironment cells. UMAP embeddings for samples from (A) GSE154109, (B) GSE185381, (C) GSE235063, (D) GSE235923 training samples, (E) GSE236923 test samples, and (F) scPCA. Unsupervised clusters produced by the Louvain algorithm (FindClusters function in Seurat) were scored for patient specificity, cytogenetic group specificity, and the proportion of healthy control cells present in the cluster. Clusters were labeled “likely malignant” if >= 70% specific to a single patient or cytogenetic group, and had <= 2% healthy control cells. Clusters meeting one of the two criteria were labeled “ambiguous”, and cells meeting neither were labeled “likely microenvironment”. Clusters with >=80% lymphoid or erythroid cells were labeled “lymphoid/erythroid predominant”.

**Figure S2:**
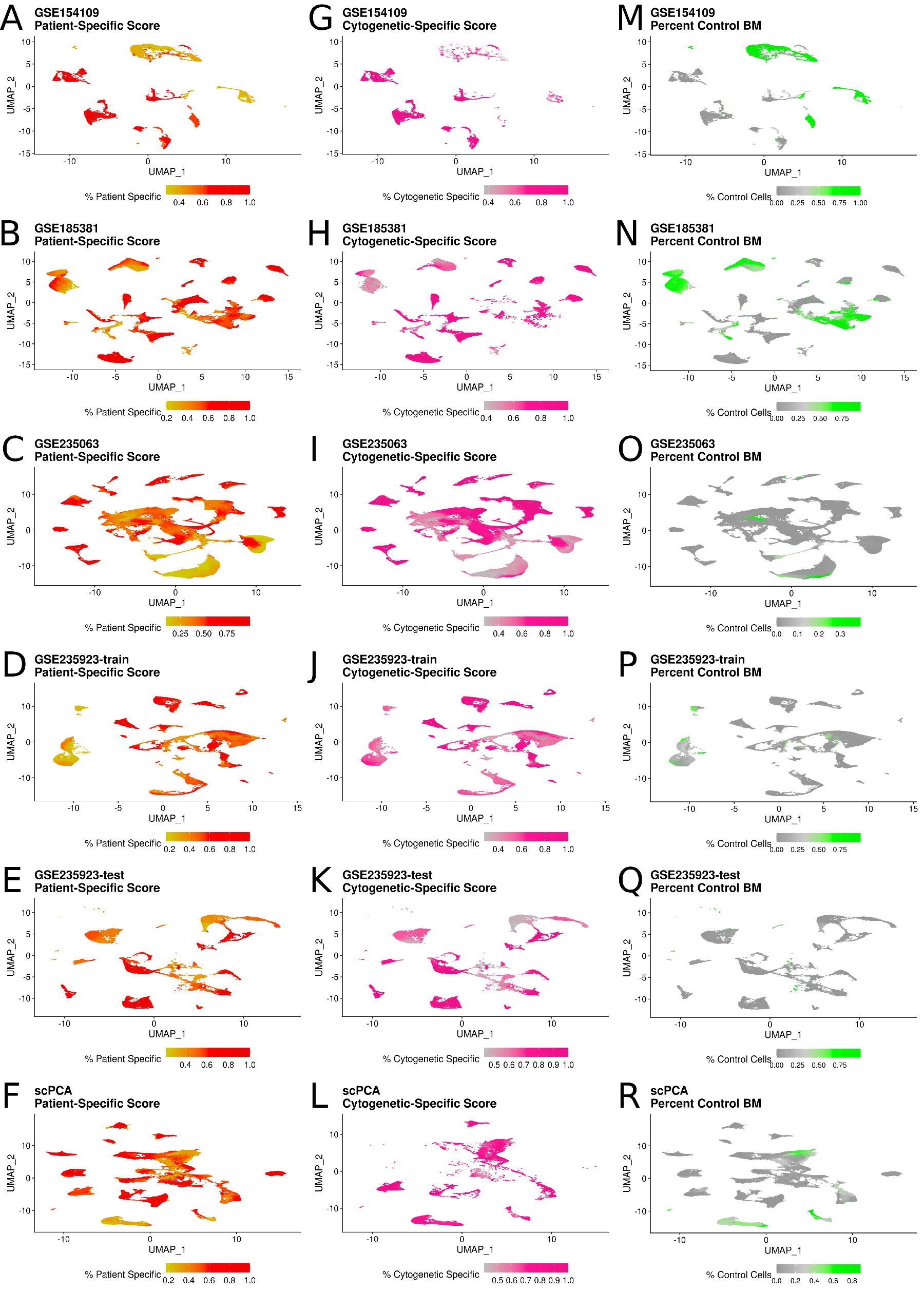
Dataset-wise criteria used to annotate clusters as malignant or microenvironment. (A-F) UMAP embeddings for each dataset with a color scale for patient-specific occupancy score (red: high patient occupancy score, yellow: low patient occupancy score). Patient-specific occupancy score was calculated by dividing the number of cells in a cluster from a single patient by the total number of cells in that cluster. Clusters were assigned the value of the patient with the highest percentage. (G-L) UMAP embeddings for each dataset with a color scale for cytogenetic-specific occupancy score (pink: high cytogenetic occupancy score, gray: low cytogenetic occupancy score). As with patient-specific occupancy, values were calculated by dividing the number of cells from a single cytogenetic group by the total cells in the cluster and assigning the cluster the highest value of all groups. (M-R) UMAP embeddings for each dataset with a color scale for the percent of cells from healthy control samples (green: high percent of control cells, gray: low percent of control cells). For each cluster, the number of cells from control samples was divided by the total number of cells in the cluster.

**Figure S3:**
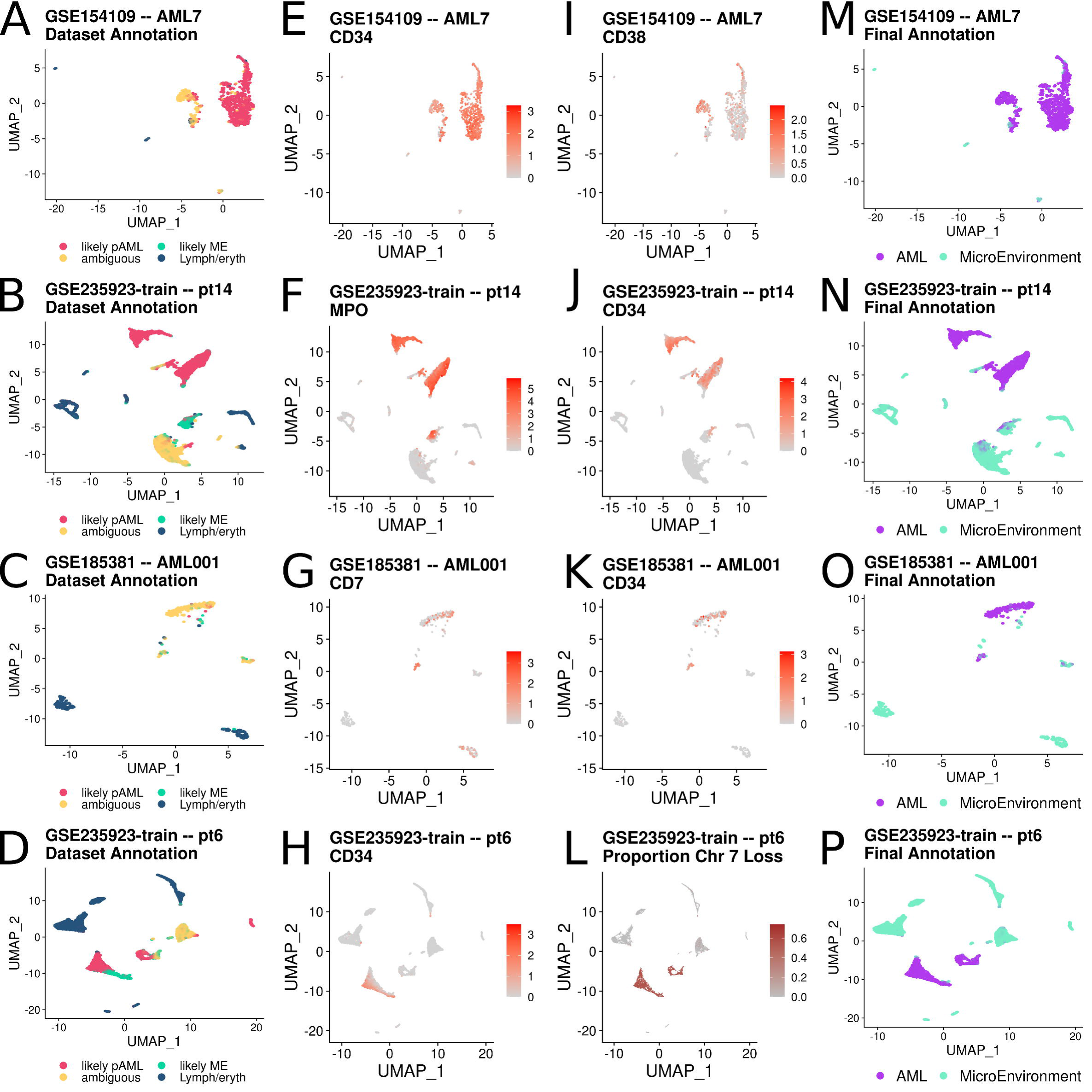
Representative plots of patient-wise malignant cell annotation. (A-D) UMAP embeddings for the cells from single patients colored by the initial malignant annotation assigned at the dataset level (as shown in Fig S1-2). (E-F) Feature plots showing gene expression for blast markers. Points are colored by normalized expression with red for high expression and gray for low. (G) Feature plot showing the proportion of chromosome 7 loss in myeloid cells from patient 6 from GSE235923. Patient 6 had chromosome 7 deletion detected clinically, so inferCNV was used to detect which myeloid cells were inferred to have chromosome 7 loss with lymphoid and erythroid cells as the reference set. (M-P) UMAP embeddings with color indicating malignant (purple) or microenvironment (turquoise) annotation. Each row illustrates how dataset-level annotations were updated to finalized malignant or microenvironment annotations. Row 1 (AML7) shows how the blast marker gene expression was used to identify two groups of pAML cells (one predominantly “likely malignant”, one “ambiguous”). Row 2 (pt14) illustrates how blast markers which are also common myeloid markers were applied for annotation. *MPO* (myeloperoxidase) is a common monocyte marker. Three groups of cells were *MPO*^+^, two of which were “likely malignant” and one that was “likely microenvironment”. In this case, the two “likely malignant” groups were also *CD34*^+^, and were annotated “malignant”. Conversely, the “likely microenvironment” group was *CD34*^-^ and was annotated “microenvironment” because it likely represents a group of non-malignant monocytes. Row 3 (AML001) illustrates a similar logic for myeloid cells. AML001 was found to have *CD7* as a blast marker by flow cytometry. *CD7* is a widely expressed gene found also in lymphoid cells. “Ambiguous” myeloid cells that were *CD7*^+^ were annotated “malignant”, but lymphoid cells that were *CD7*^+^ were annotated “microenvironment”. Row 4 illustrates the use of inferCNV to identify malignant cells in patient 6. Patient 6 had several myeloid groups of cells with mixed dataset-level annotations. One group of cells was *CD34*^+^ (detected by flow cytometry), and two groups of cells had chromosome 7 loss by inferCNV. Cells from both CNV^+^ myeloid groups were annotated “malignant”, and CNV^-^, blast marker^-^ cells were annotated microenvironment.

**Figure S4:**
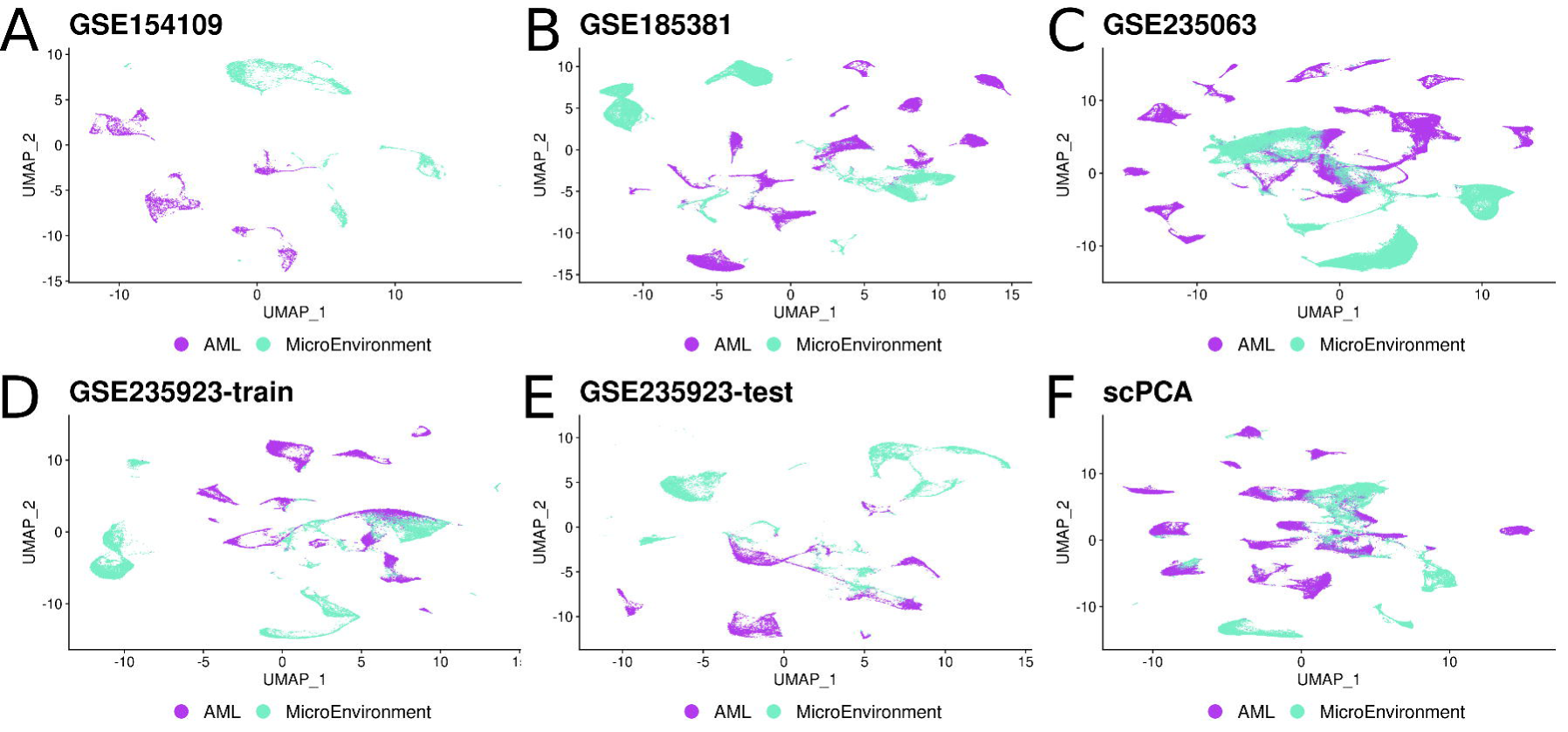
Final annotation of malignant pAML cells and non-malignant microenvironment cells. (A-F) UMAP embeddings for each dataset, points are colored by final annotation as malignant (AML) or non-malignant (microenvironment). The color indicates annotation with purple for AML and teal for microenvironment.

**Figure S5:**
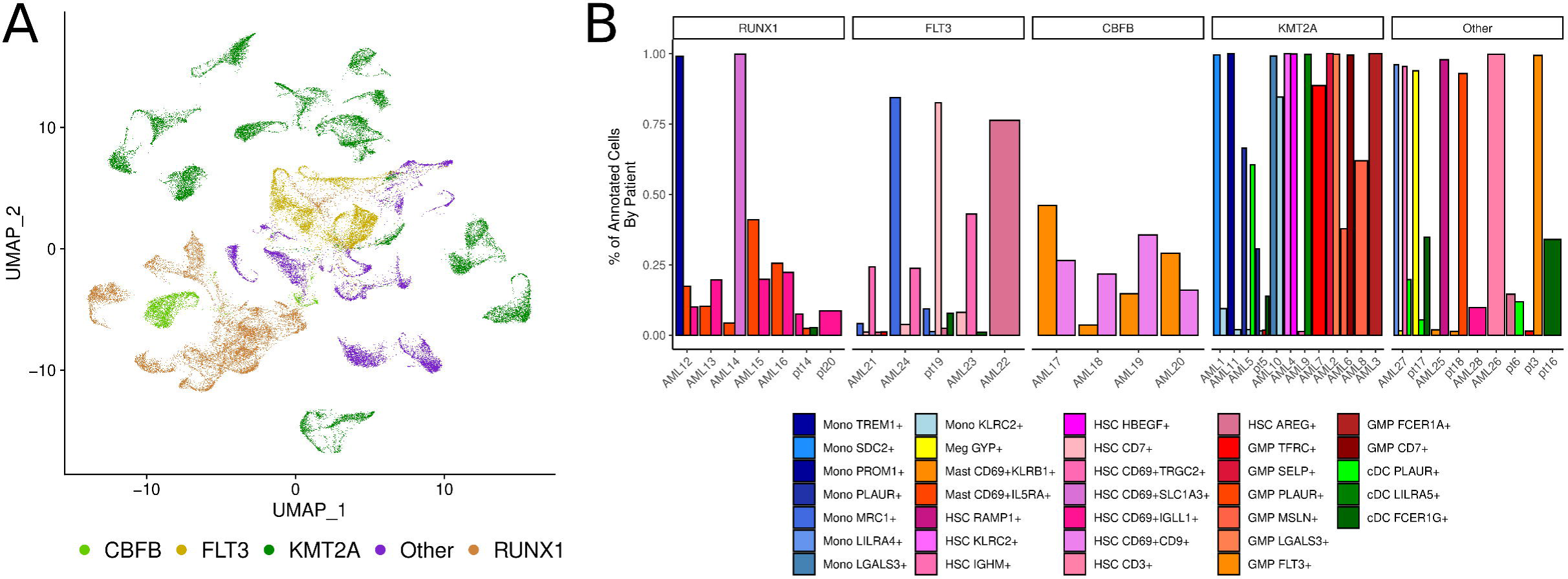
Patient-specificity of malignant subtypes across cytogenetic categories. (A) UMAP embedding for malignant cells from 37 patients with matched diagnosis, end-of-induction, and relapse samples. Cells are colored by cytogenetic group (lime: CBFB-MYH11, yellow: FLT3-ITD, green: KMT2A rearranged, brown: RUNX1-RUNX1T1, purple: other). (B) Bar chart showing the patient-specificity of malignant subtypes. Bar height indicates the percentage of cells from a subtype that were derived from each patient. Subtypes in the KMT2A and “other” categories were >90% specific to a single patient in 17 out of 21 cases as compared to 2 out of 16 from RUNX1, FLT3, and CBFB. This was interpreted to indicate that RUNX1, FLT3, and CBFB pAML subtypes are conserved between patients to a greater extent than KMT2A and “other” groups.

**Figure S6:**
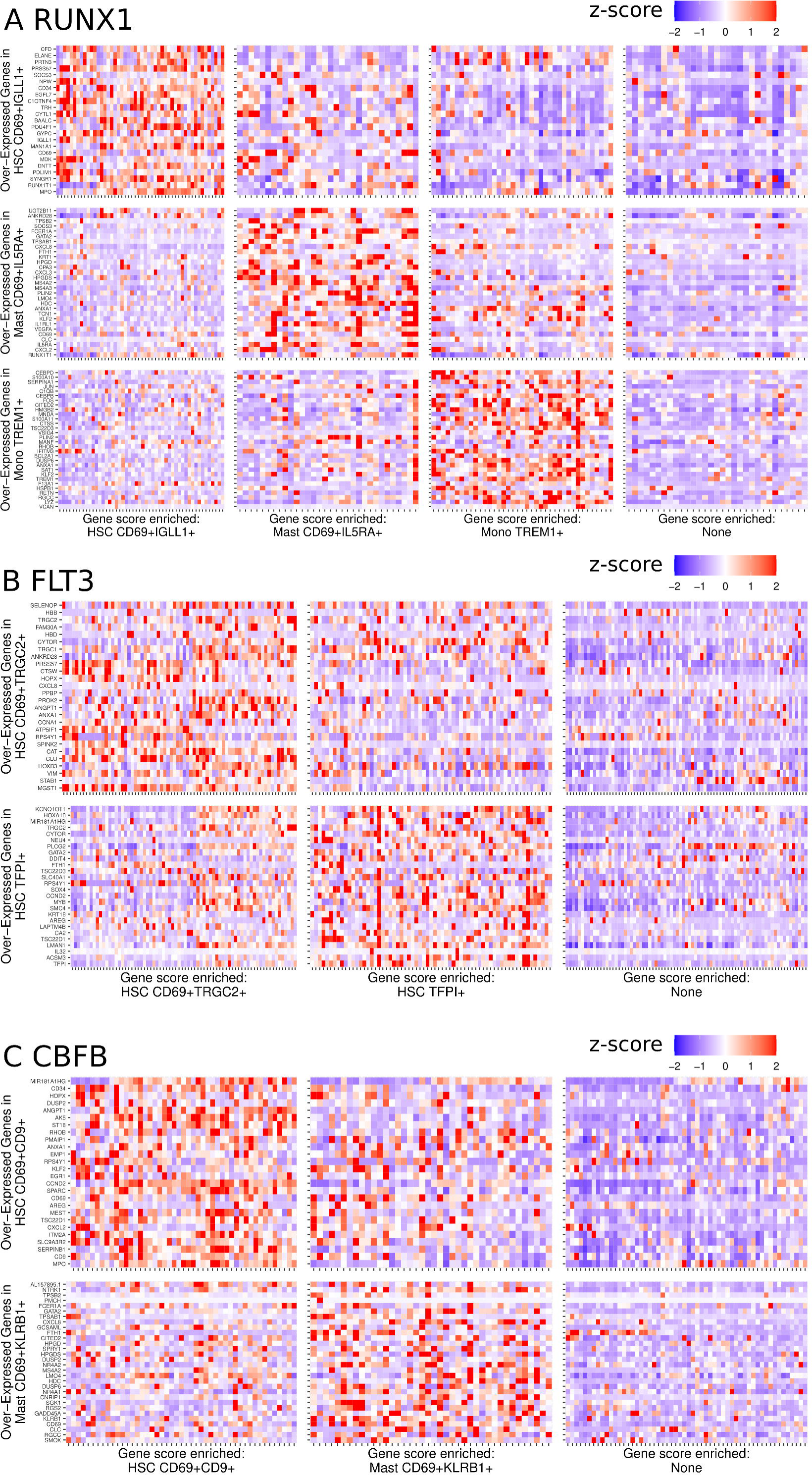
Deconvolution of TARGET bulk RNAseq data with pediatric acute myeloid leukemia (pAML) subtype gene signatures. Heatmap for (A) RUNX1-RUNX1T1, (B) FLT3-ITD, and (C) CBFB-MYH11 samples. Genes were selected based on the top 30 genes by log_2_ fold change with log_2_ fold change > 0.5, adjusted p < 0.05. Significance was tested by Wilcoxon rank sum test for each subtype (shown on the Y-axis). Gene signature scores were calculated from the mean normalized z-score for each gene in the signature. Each sample was scored for all gene signatures from its corresponding cytogenetic group and then grouped into the subtype with the highest signature score (or none of all gene signature scores were negative). The color indicates normalized, z-scaled gene expression (red: high, blue: low) for highly expressed genes in the high-risk, treatment-resistant subtypes of pAML identified in scRNAseq. TARGET: Therapeutically Application Research to Generate Effective Treatments, pAML: pediatric acute myeloid leukemia, scRNAseq: single-cell RNA sequencing.

**Figure S7:**
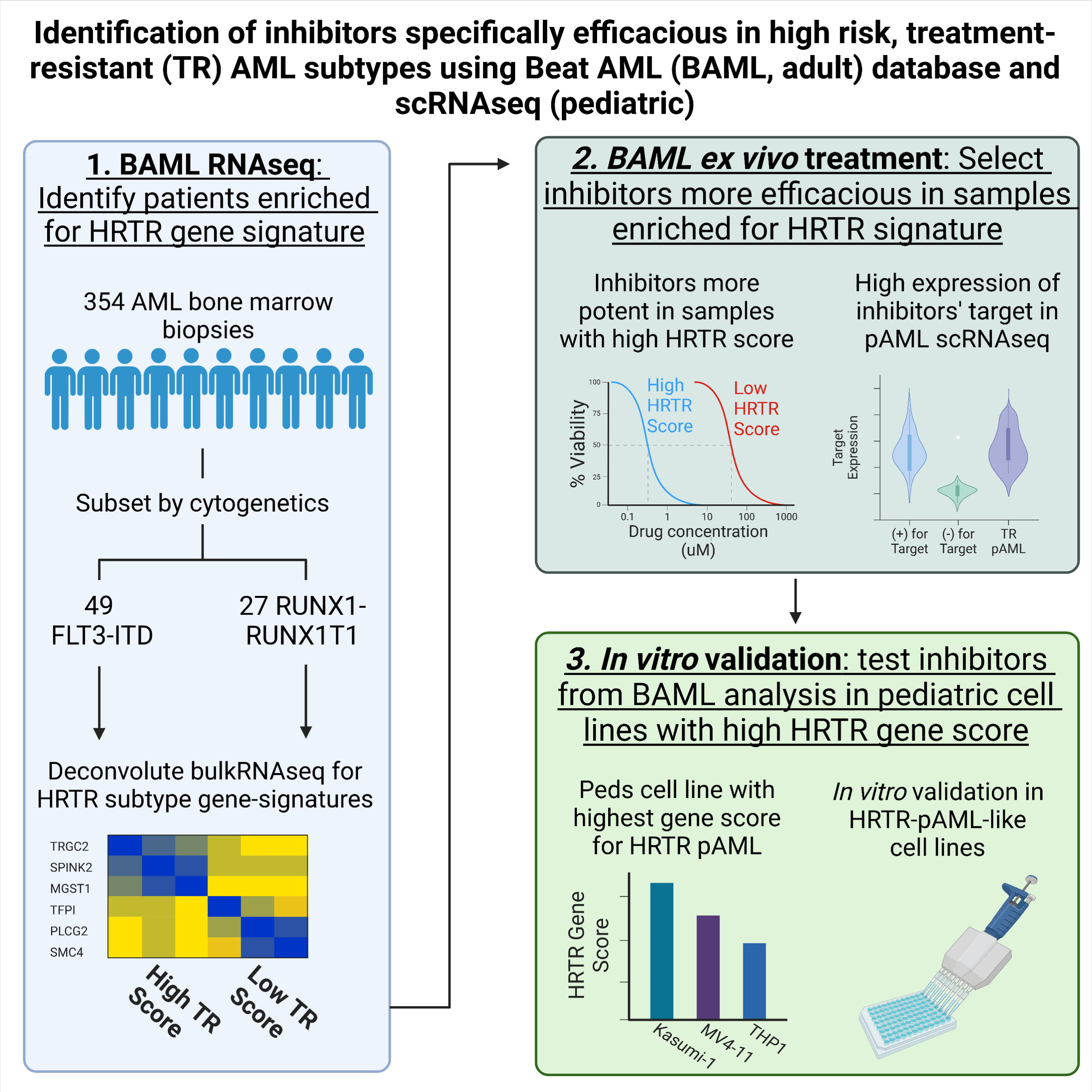
Schematic for identification of pharmaceutical inhibitors specifically efficacious in high-risk, treatment-resistant (HTRT) pediatric acute myeloid leukemia (pAML). Gene signatures constructed from the top 30 highly expressed genes (log_2_ fold change > 0.5, adjust p value < 0.05) in HRTR pAML subtypes (scRNAseq) were used to deconvolute bulk RNA sequencing samples in the Beat AML database. Gene signature scores were calculated by the mean normalized, z-scaled gene expression for all genes in the gene signature. Samples were considered “high” or “low” for the HRTR gene signature based on a signature score greater or less than the mean for the cytogenetic group (e.g. all FLT3 samples for a FLT3 subtype of HRTR pAML). The percent viability of pAML cells after e*x vivo* treatment was compared between samples from the high and low HRTR groups to identify inhibitors with higher potency in samples with similar gene expression to HRTR pAML subtypes. Given that inhibitors were identified in an adult AML database, we used the expression of genes or pathways targeted by the inhibitors with relatively high or low potency to provide pediatric evidence to support the *ex vivo* findings. Finally, pediatric cell lines with high expression of HRTR pAML gene signatures were used for *in vitro* validation of the inhibitors identified in the Beat AML analysis.

**Table S1:** Single-cell RNA sequencing datasets from pediatric acute myeloid leukemia samples used in this project.

**Table S2:** Clinical metadata for single-cell RNA sequencing samples from pediatric acute myeloid leukemia.

**Table S3:** Genes used for cell type annotation in single-cell RNA sequencing datasets.

**Table S4:** Metrics used malignant vs microenvironment annotation of single-cell RNA sequencing datasets.

**Table S5:** Marker genes for *CD69*^+^*IGLL1*^+^ hematopoietic stem cell-like pediatric acute myeloid leukemia cells versus other pediatric acute myeloid leukemia cells.

**Table S6:** Marker genes for *CD69*^+^*TRGC2*^+^ hematopoietic stem cell-like pediatric acute myeloid leukemia cells versus other pediatric acute myeloid leukemia cells.

**Table S7:** Marker genes for mast cell-like pediatric acute myeloid leukemia cells versus other pediatric acute myeloid leukemia cells.

