## Supplemental Methods for "Single-cell analysis of pediatric acute myeloid leukemia samples uncovers treatment-resistant stem and mast cells"

**Materials and Methods (Supplemental)**

*Single-cell RNA sequencing datasets, processing, and cell type identification*

ScRNA-seq datasets were procured from the Gene Expression Omnibus (NCBI GEO) (GSE235923, GSE235063, GSE154109, GSE185381) and ScPCA (https://scpca.alexslemonade.org/) databases, including datasets recently published from our lab^1^ (**Supplemental Table 1**). Low-quality cells were filtered out by removing cells with <250 features, >6,000 features, and >20% mtDNA transcripts. Potential doublets were marked patient-wise using DoubletFinderV3^2^. Clusters with >50% cells marked as potential doublets and expression of canonical genes from at least 2 different cell types (e.g. *HBB* and *CD14*, see **Supplemental Table 3**) were removed from further analysis. After low quality and doublet cell removal, each dataset was independently processed using the Seurat package^3^. Raw counts per cell were log-normalized using the NormalizeData function in the Seurat package. To address the variation in expression levels of different genes, we applied gene-wise scaling using the ScaleData function in Seurat^3^. Normalized, and scaled expression values for the top 2000 variable genes were used for principal component analysis (PCA). Next, to identify groups of cells with similar gene expression, we constructed a K-nearest neighbor (KNN) graph based on the Euclidean distance of the first 10 principal components using the FindNeighbors function in Seurat. Then the Louvain algorithm was applied to the KNN graph to group cells into clusters of cells with similar expression profiles. To visualize cells in a lower dimensional representation we performed uniform manifold approximation and projection (UMAP) using the RunUMAP function in Seurat. Batch correction was performed to remove dataset-specific effects using the integration anchors approach. Cell types were annotated using canonical cell type-specific marker gene expression (**Supplemental Table 3**)^4^.

*Malignant blast cell identification – Step 1: Dataset-wise annotation*

To differentiate malignant pAML blast cells from healthy myeloid cells, we applied a two-step annotation process (**Fig 2D**). First, clusters of cells identified were analyzed data-set-wise for the percentage of cells that were derived from a single patient, the percentage of cells that were derived from a single cytogenetic group, and the percentage of cells from healthy control BM samples. Datasets that did not have healthy control samples (GSE235923, GSE235063) were merged with the four healthy control samples from the GSE154109 dataset before data processing and normalization. Then dataset-wise parameters were calculated on each cluster.

1.a) Patient-specific occupancy scoring: In each cluster, the proportion of cells from individual patients was quantified. Each cluster was then assigned the score corresponding to the patient with the highest cell count (i.e. if a cluster was 99% derived from patient A and 1% from patient B, that cluster was assigned a score of 99%). Clusters were considered patient-specific based on a cut-point of >=70% as described previously^5^.

1.b) Cytogenetic group-specific occupancy scoring: As with patient-specific occupancy scoring, in each cluster the proportion of cells from individual cytogenetics groups was quantified. Clusters were considered specific to a cytogenetic group based on a cut point of >=70%, consistent with patient-specific scoring described in the previous paragraph. In datasets with larger patient counts (e.g. GSE235063, see **Supplemental Table 1**) cytogenetic scoring enabled the identification of malignant cells that were found in multiple patients from the same cytogenetic background. For example, a *CD34*^+^ cluster that was 99% derived from four patients with RUNX1-RUNX1T1 translocation would score low for patient specificity, but high for cytogenetic specificity. In datasets where each patient had a different cytogenetic background (e.g. GSE154109) patient scoring and cytogenetic scoring were equivalent. Patients without cytogenetic information were not included in this analysis.

2) Control scoring: For each cluster, the percentage of cells derived from healthy BM samples was calculated and considered depleted for healthy control cells based on a threshold of <2%.

Based on these criteria, clusters were given a putative malignant or microenvironment label. Clusters with <2% control cells and >=70% patient- or cytogenetic-specific scores were labeled “likely malignant”. Clusters with >2% control cells and < 70% patient- and cytogenetic-specific scores were labeled “likely microenvironment”. Clusters meeting one of the two criteria were labeled as “ambiguous”. Lastly, clusters that were >=80% comprised of lymphoid or erythroid cells were labeled “lymphoid/erythroid predominant” regardless of the two dataset-level criteria since malignant cells were expected only in the myeloid compartment. With established annotations from the dataset-wise analysis, we next analyzed each patient independently to finalize malignant and microenvironment annotation.

*Malignant blast cell identification – Step 2: Patient-wise annotation*

Next, for patient-wise malignant annotation strategies (blast marker gene detection, CNV detection) the feature barcode matrices for individual patients were separately normalized, scaled, dimensionally reduced with PCA, clustered with KNN and Louvain algorithms, and visualized using UMAP. Four of the six datasets had available clinical flow cytometry metadata (**Supplemental Table 2**). In these datasets, patients were individually analyzed for high expression of blast markers identified by flow cytometry. Cells were annotated “AML” if expressing genes for blast markers, except for cases where the dataset annotation was "likely microenvironment” and the blast cell annotation marker is also a marker of normal myeloid cells in the bone marrow. For example, given a patient with *CD14* as a blast marker by flow cytometry, the *CD14*^+^ cells that were annotated “likely malignant” were given a final “malignant” annotation. Conversely, *CD14*^+^ cells that were annotated “likely microenvironment” were given a final annotation of “microenvironment”. For the two datasets without flow cytometry metadata, we used markers that are highly over-expressed by malignant blast cells in the BM (HSC: CD34, MSI2, GMP: AZU1, Mast: TPSAB1, FCER1A). For the second patient-wise annotation strategy, fourteen patients had clinical CNV meta-data (**Supplemental Table 2**). These patients were analyzed by InferCNV^6^ using non-myeloid cells as the reference set. Cells with the expected chromosome gain or loss (e.g. chromosome 7 loss in a patient with chromosome 7 deletion) by inferCNV were annotated “AML”. Finally, “AML” and “microenvironment” cell labels were added to the final cellular annotation meta-data for all downstream analyses.

*Correlative validation of Malignant AML cell annotation*

Malignant annotations were verified in two ways: 1) The percentage of cells annotated as malignant in scRNAseq was correlated with the blast percentage identified by clinical flow cytometry using Pearson correlation analysis. The correlation was considered significant if the p-value < 0.05. 2) Malignant annotations were compared to previously published malignant annotations from GSE185381 for verifying consistency.

*Identification of high-risk, treatment-resistant pAML subtypes*

A broad goal of this project was to identify subtypes of pAML that promote relapse within existing cytogenetic categories. To do so we analyzed the malignant cells from 37 patients who had at least two matched samples from diagnosis (DX), end-of-induction (EOI), and/or relapse (REL). To summarize the process (elaborated in subsequent sections): We first identified subtypes of pAML by differential expression of genes that code for cell surface proteins (e.g. HSC-like *CD69*^+^ pAML). With these pAML subtypes, three criteria were used to identify “high risk, treatment-resistant” (HRTR) pAML: 1) pAML subtypes were considered relapse-associated if present as a greater proportion of malignant cells in patients who experienced relapse as compared to patients with sustained remission. 2) pAML subtypes were considered treatment-resistant if they accounted for a greater proportion of malignant cells at EOI as compared to DX. 3) pAML subtypes were considered high risk if they expressed a gene signature associated with shortened overall survival in the Therapeutically Applicable Research to Generate Effective Treatments (TARGET) data. PAML subtypes meeting all three criteria were considered HRTR.

*Annotation of pAML subtypes in matched diagnosis, end-of-induction, and relapse samples*

To characterize pAML that is associated with relapse, we first sought to define subtypes of pAML present in the 37 patients with matched diagnosis (DX), end-of-induction (EOI), and relapse (REL) samples (36 DX, 34 at EOI, 27 at REL). Feature barcode matrices for patients with matched samples were merged, batch corrected using Seurat integration anchors, normalized, scaled, dimensionally reduced by PCA, clustered using KNN and the Louvain algorithms, and visualized using UMAP. To examine specifically the pAML cells, microenvironment cells (e.g. T cells, B cells, non-malignant myeloid cells) were removed from the analysis.

To annotate subtypes of pAML, we identified genes for cell surface markers that were differentially expressed (log_2_ fold change > 0.5, adjusted p < 0.05) in pAML clusters. The Louvain algorithm (FindClusters function) identified 60 clusters of pAML. Differential expression was performed using the FindMarkers function with the Wilcoxon rank sum test between each cluster vs. all other clusters. Differentially expressed genes were subset to those coding for cell surface proteins using the gene list from GO:0009986, the “Cell Surface” list in the HUGO gene nomenclature committee, and manual identification in the NCBI gene reference database. Clusters with similar top differentially expressed genes corresponding to cell surface proteins were annotated as the same subtype. For example, two HSC-like clusters with high expression of *CD69* would be labeled “HSC *CD69*^+^”. Canonical cell-type annotation genes were not used for subtype labeling. For example, an HSC-like pAML cluster that differentially expressed *CD34* would not be labeled *CD34*^+^ because *CD34* was used as a marker gene for the identification of HSCs. Genes that were expressed in >50% of all malignant cells (e.g. heat shock proteins, annexins, human leukocyte antigens) were not used for annotation except *CD69* which has previously been described as a leukemic stem cell marker^7^.

*Identification of relapse-enriched subtypes of pAML within cytogenetic categories*

Identification of pAML subtypes associated with relapse was carried out by comparing the pAML subtypes that were present in patients who experienced relapse vs those who achieved sustained remission (**Supplemental Table 2**). The 37 samples with matched time points were predominantly from patients who experienced relapse (27/37). To improve the balance of patients who experienced relapse vs sustained remission we included samples from patients who had only diagnosis samples and outcomes metadata (relapse/remission). Feature barcode matrices for matched and un-matched samples with clinical outcomes were merged, batched corrected using the Seurat integration anchors approach, normalized, scaled, dimensionally reduced by PCA, clustered using KNN and the Louvain algorithm, and visualized using UMAP. Analysis of samples from each cytogenetic category (RUNX1, FLT3, CBFB) was conducted independently. Cells were annotated based on differentially expressed cell surface genes using the FindMarkers function in Seurat with the Wilcoxon rank sum test as described for matched samples. The proportion of pAML subtypes in relapse/remission samples was calculated by dividing the number of cells from a pAML subtype by the total number of malignant cells with the same cytogenetic and outcome group. Log_2_ fold proportion was calculated by dividing the proportion of a subtype in relapse samples by the proportion of a subtype in remission samples. For example, the proportion of *CD69*^+^*IGLL1*^+^ HSC-like cells in RUNX1 relapse samples was divided by the proportion of *CD69*^+^*IGLL1*^+^ HSC-like cells in RUNX1 remission samples. To reduce the false positives, the subtypes analysis was performed on subtypes with at least 250 cells in at least 1 patient and at least 1 outcome group. Subtypes with a positive log_2_ fold proportion in patients who experienced relapse vs sustained remission were considered relapse enriched.

*Identification of treatment-resistant subtypes of pAML within cytogenetic categories*

To further identify which subtypes of pAML were resistant to treatment we quantified the extent to which cells persisted after treatment within cytogenetic categories. pAML cells were grouped by cytogenetic category. The proportion of subtypes at each time point was calculated by dividing the number of cells from a given subtype (e.g. *CD69*^+^*IGLL1*^+^ HSC-like cells from RUNX1 patients at diagnosis) by the total number of cells in that cytogenetic group at that time point. To restrict the comparison to abundant subtypes, subtypes were compared within cytogenetic categories for all subtypes with at least 250 cells in at least one patient and at least one time point. AML subtypes were considered treatment-resistant if that subtype had a proportional increase in cell abundance at EOI relative to DX.

*Generation of gene signatures for relapse-associated pAML subtypes*

To determine if the transcriptome profiles of relapse-associated pAML subtypes were associated with outcomes in a larger cohort of patients, we generated subtype gene signatures. The signatures were generated by marker analysis comparing the target subtype with all other subtypes using the FindMarkers function with the Wilcoxon rank-sum test for significance testing. The gene signature for each subtype contained genes with log_2_ fold change > 0.5 and adjusted p < 0.05. If the analysis identified >30 genes significantly different, the top 30 genes based on FC were selected for inclusion in the gene signature. These pAML subtype gene signatures were subsequently used for the deconvolution of the TARGET pAML bulk RNA sequencing data.

*Identification of high-risk subtypes of pAML by TARGET deconvolution and risk analysis*

To explore survival outcomes of relapse-associated pAML subtypes in a larger cohort, data were downloaded from the TARGET pAML initiative (<https://www.cancer.gov/ccg/research/genome-sequencing/target>). Bulk RNA sequencing samples were subset to BM biopsies collected at the time of disease diagnosis from patients on the AAML03P1, AAML0531, and AAML1031 protocols for relative consistency of treatment. For analysis of each relapse-associated pAML subtype, TARGET samples were subset to those that had the same cytogenetic mutation as the pAML subtype being analyzed (e.g. subset to RUNX1-RUNX1T1 samples in TARGET for a pAML subtypes found in RUNX1-RUNX1T1 patients in scRNAseq). Samples were excluded if they had co-occurring mutations from the other cytogenetic categories (e.g. for a RUNX1-RUNX1T1 patient, exclude for co-occurring FLT3-ITD, CBFB-MYH11, or KMT2A rearrangement), X chromosome deletion and KIT mutation which were both not well represented in our cohort, or had cytogenetic metadata marked "Unknown". The bulk RNA-seq data was normalized using counts-per-million and z-scaling. To calculate a gene signature score for each pAML subtype from a given cytogenetic category, normalized z-scores for each gene were averaged across all genes from each gene signature. For example, RUNX1 had three relapse-associated subtypes, so RUNX1 bulkRNAseq samples in TARGET were scored for all three signatures. Samples were labeled by the highest gene signature score with a value > 0 (e.g. if the highest gene signature score was for *CD69*^+^*IGLL1*^+^ HSC-like pAML, it was labeled “HSC *CD69*^+^*IGLL1*^+^ high”). Samples with negative mean z-scores for all gene signatures from that cytogenetic category were labeled enriched for “none” of the pAML subtypes. To determine if there was a difference in overall survival between samples enriched for different pAML subtypes, survival analysis was performed using the Survival R package with Cox proportional hazards model for significance testing^8^. The subtypes were considered significantly associated with poor outcomes if the hazard ratio was > 1.0 and p < 0.05.

In summary, pAML subtypes that met all three of the following criteria were classified as "high risk, treatment-resistant" (HRTR) pAML: 1) Proportionally enriched in patients who experienced relapse versus those who sustained remission, 2) Proportionally enriched at the end of induction (EOI) compared to diagnosis, and 3) Expressed a gene signature associated with a statistically significant hazard ratio greater than 1.0 in corresponding bulk RNA-seq data from the TARGET initiative.

*Pathway and gene-set enrichment analysis for high-risk, treatment-resistant pAML subtypes*

To understand the biological functions driving HRTR pAML, we performed pathway and gene-set enrichment analysis. Pathway analysis was performed on over-expressed genes (i.e. positive fold change value, adjusted p-value < 0.05) using the Kyoto Encyclopedia of Genes and Genomes (2021), Reactome, and GO_Biological Process (2023) using the EnrichR package^9–12^. Pathways with multiple test-corrected p-value <0.05 were considered significant. Further gene set enrichment analysis for selected pathways relevant to the inhibitors identified as relatively high or low efficacy in HRTR pAML (WP_PROTEASOME_DEGRADATION, REACTOME_MTOR_SIGNALING) was performed using the fgsea package^13^ on the rank-ordered list of genes for each subtype and considered significant if p-value <0.05.

*Identification of drugs targeted to patients with elevated expression of high-risk, treatment-resistant gene signatures – Overview*

After identifying HRTR pAML subtypes, we next sought to identify inhibitors specifically efficacious for patients enriched with these cells. To achieve this, we used the adult Beat AML bulk RNA sequencing and *ex vivo* treatment data in combination with our pAML scRNAseq data (**Supplemental Figure 7**)^14^. First, bulk RNA sequencing samples from the Beat AML dataset were deconvoluted to identify samples with high (and low) enrichment for HRTR pAML gene signatures. Then, to identify inhibitors with higher efficacy in samples enriched for HRTR gene signatures, we compared cell viability after *ex vivo* treatment between samples from the “high” and “low” gene signature groups. Since Beat AML is an adult database, we provided supporting pediatric evidence by analyzing our scRNAseq data for differential expression of targets, pathways, or sensitivity genes of the inhibitors with differential efficacy. Finally, inhibitors with high or low efficacy and supporting scRNAseq evidence were validated *in vitro* using pediatric cell lines.

*Identification of bulk RNA sequencing samples enriched for high-risk, treatment-resistant gene signatures in the Beat AML database*

The first step in identifying inhibitors efficacious against HRTR pAML was to identify samples in the Beat AML bulk RNA sequencing data set that were enriched for HRTR pAML gene signatures. Bulk RNA sequencing samples were subset to BM biopsies from the same cytogenetic background as the HRTR pAML subtype (e.g. RUNX1 samples for *CD69*^+^ *IGLL1*^+^ HSC-like pAML subtype). Each bulk RNA sequencing sample was scored for HRTR pAML gene signature by the average z-scaled normalized expression for all genes in the signature. Samples were considered enriched (“high”) for the HRTR gene signature if the signature score was greater than the average signature scores for all samples within the cytogenetic group and vice versa for not enriched (“low”). With samples labeled as “high” or “low” enrichment for HRTR pAML gene signatures, the next step was to compare cell viability after *ex vivo* treatment.

*Identification of inhibitors with higher efficacy in samples enriched for high-risk, treatment-resistant gene signatures in the Beat AML ex vivo treatment data*

With samples stratified into “high” or “low” HRTR pAML groups, we next aimed to identify inhibitors with greater efficacy in *ex vivo* treatment samples enriched for HRTR pAML gene signatures. The Beat AML *ex vivo* treatment data contains cell viability normalized to untreated control wells for inhibitor concentrations from 0.00137 to 10.0 µM. For these methods, we define an “assay” as the experiment including all concentrations from one sample and inhibitor (e.g. all wells from 0.00137 to 10 uM for Sample-2440 treated with bortezomib). To minimize confounding due to *ex vivo* treatment duration, assays that were treated for less than 20 hours were excluded from the analysis. Additionally, multiple criteria were used to filter out low-quality assays: 1) Assays with higher cell viability at the highest dose (10 µM) as compared to the lowest dose (0.0137 µM), 2) Assays with viability showing an increase of >50% from between two incrementally increasing doses of inhibitor, 3) Assays with normalized viability of 0 for all concentrations, 4) Assays with normalized viability greater than 100% for all concentrations, 5) Individual wells with viability >200%. After filtering to high-quality *ex* vivo assays, viability values were compared between HRTR high and low samples. Normalized viability versus inhibitor concentration was modeled by a 5-parameter log-logistic model for the HRTR gene signature high and low groups independently. Using the logistic model, 50% inhibitor concentrations (IC_50_) were estimated. Then, to provide a metric of relative efficacy, log_2_ fold potency (L_2_FP) was calculated by log_2_(IC_50-high_/ IC_50-low_). Statistical significance was tested by two-way ANOVA (Normalized viability ~ gene signature group + well concentration + gene signature group:well concentration). P value for gene signature group effects was used to define statistical significance in efficacy. Inhibitors were considered differentially efficacious by an absolute L_2_FP >= 1.0 and p < 0.05. Inhibitors with absolute L_2_FP >= 1.0 and p < 0.05 were next analyzed for supporting transcriptomic evidence in our pediatric scRNAseq data.

*Supporting transcriptomic evidence for inhibitors identified as highly efficacious in Beat AML*

Given that an adult *ex vivo* treatment dataset was used to predict efficacy for subtypes of pAML, we next aimed to find supporting evidence in our pediatric scRNAseq data. Specifically, we analyzed the HRTR subtypes of pAML for enrichment of genes or pathways that either are targets of the inhibitors identified or associated with sensitivity to the inhibitors. For example, if the JAK inhibitor ruxolitinib was found to have higher efficacy in Beat AML samples enriched for HRTR pAML (L_2_FP >= 1.0, p < 0.05), then 1) High expression of JAK2, 2) Enrichment for the “JAK signaling” pathway, or 3) High expression of a gene previously reported as a predictor of sensitivity would be considered supporting evidence. For assessing genes that code for an inhibitor’s target, we grouped samples as + or – based on whether the subtype differentially expressed the corresponding gene. As an example for the ANGPT1 gene, all pAML subtypes that differentially expressed ANGPT1 (L_2_FC > 0.25, adjusted p < 0.05) were labeled ANGPT1^+^, and other subtypes were labeled ANGPT1^–^ (typically with 0 counts in >80% of cells). For genes published as predictors of sensitivity to a given inhibitor, the same process was applied except that subtypes were labeled “high” or “low” as sensitivity genes typically had non-0 count detection in >50% of cells. Finally, gene-set enrichment analysis was used to determine if there was enrichment for pathways targeted by inhibitors with absolute L_2_FP >= 1.0, p <0.05. For example, Bortezomib was found to be more efficacious in samples that were enriched for the *CD69*^+^ *IGLL1*^+^ HSC-like gene signature, so fgsea was used to test for enrichment of the Wiki Pathways Proteasome Degradation pathway (cut point of p < 0.05). Inhibitors with an absolute L_2_FP >= 1.0, p < 0.05 by ANOVA, and transcriptomic support for the efficacy observed in Beat AML were selected for *in vitro* validation.

*In vitro validation of targeted inhibitors for high-risk, treatment-resistant pAML*

To validate the inhibitors predicted to be relatively high or low efficacy in HRTR pAML, we performed *in vitro* drug sensitivity testing of pAML cell lines. Cell lines were selected based on enrichment for HRTR pAML subtype gene scores and the presence of the cytogenetic mutation corresponding to the subtype of pAML. The GSE59808 dataset has microarray data for several pediatric and adult cell lines. Normalized expression values were z-scaled gene-wise to equally weigh all genes in the gene signatures. Gene signature scores for *CD69*^+^ *IGLL1*^+^ HSC-like pAML (RUNX1) and *CD69*^+^ *TRGC2*^+^ HSC-like pAML (FLT3) were calculated based on the average normalized z-scaled expression for all signature genes. Kasumi-1 (RUNX1) and MV4-11 (FLT3) were found to have the highest gene score for *CD69*^+^ *IGLL1*^+^ HSC-like pAML (RUNX1) and *CD69*^+^ *TRGC2*^+^ HSC-like pAML (FLT3) respectively.

To test *in vitro* drug sensitivity, Kasumi-1 and MV4-11 cells were cultured to 2500 count per well with 90% viability in a total volume of 50 µL cell culture complete media. Cells are treated with inhibitors at incremental concentrations from 0-118uM (except in H-89 and nilotinib which were extended up to 1000 µM due to low efficacy). The inhibitors tested included bortezomib (17324-69-7), ponatinib (943319-70-8), venetoclax (1257044-40-8), nilotinib (641571-10-0), H-89 (127243-85-0), rapamycin (53123-88-9), and PD173955 (260415-63-2), purchased from medchemexpress.com. After 72 hours, 100 µL/well of CellTiter-Glo® 2.0 (Cat#G9241, Promega) was added to each well, and cell viability was assessed by measuring the absorbance at 490 nm in a 96-well plate reader (BMG, Labteck, CLARIO Star). The doses that decreased cell viability to 50% (IC_50_) were analyzed using a nonlinear log regression (inhibitor) vs. normalized response (three parameters) with GraphPad Prism software.

*Evaluation of lymphoid subtype abundance in RUNX1 high-risk, treatment-resistant pAML*

Differential expression and pathway analysis of *CD69*^+^ *IGLL1*^+^ HSC-like pAML indicated that it may have an impact on the immune microenvironment. To examine the potential impact on the immune microenvironment, we analyzed the lymphoid cells from RUNX1 patients. All samples from RUNX1-RUNX1T1 pAML were merged, batch corrected using the Seurat integration anchors approach, normalized, scaled, dimensionally reduced by PCA, clustered using KNN and the Louvain algorithm, and visualized using UMAP. Lymphoid subtypes were identified using canonical gene expression^5^.

To compare the lymphocytes from patients enriched for *CD69*^+^ *IGLL1*^+^ HSC-like pAML versus those who were not, we identified the most abundant pAML subtype in each sample. Samples in which *CD69*^+^*IGLL1*^+^ HSC-like pAML was the malignant cell type with the highest cell count were labeled “*CD69*^+^*IGLL1*^+^”, while samples with any other pAML subtypes as the highest cell count were labeled “other”. The proportional abundance of lymphoid subtypes was compared between *CD69*^+^*IGLL1*^+^ and other patients and significance testing was performed using student’s t-test. To determine if similar trends between *CD69*+*IGLL1*+ HSC-like pAML and lymphoid subtypes were present in TARGET, we deconvoluted diagnostic bone marrow biopsy bulk RNA sequencing samples from RUNX1 patients. Using the same method as described for survival analysis, samples were scored for the *CD69*^+^*IGLL1*^+^ HSC-like gene signature by mean normalized, z-scaled expression values for the 30 genes in the gene signature. Similarly, canonical genes for T cell subsets were used as gene signatures to calculate CD4^+^ memory (*CD4, TRADD, TIMP1, IL7R, TNFRSF4, EOMES, BCL6, LEF1*) and CD8^+^ Cytotoxic (*CD8A, CD8B, CD8B2, GZMA, PRF1, FGBP2, NKG7*) signature score. Correlation between the CD69^+^IGLL1^+^ gene signature scores and the CD4^+^ memory T cell/CD8^+^ cytotoxic T cell scores were assessed using Pearson correlation with a significance cut point of 0.05. Differential expression of lymphoid subtypes between *CD69*^+^*IGLL1*^+^ and other samples (e.g. comparison of CD8^+^ cytotoxic T cells from *CD69*^+^*IGLL1*^+^ samples vs other samples) was executed using the FindMarkers function with the Wilcoxon rank-sum test for significance testing.

*Signaling analysis between pAML and T cell subtypes*

To examine the signaling between *CD69*^+^*IGLL1*^+^ HSC-like pAML and T cells we used CellChat analysis^15^. Given that the lymphoid subtypes with significant differences in abundance were in the T cell compartment, CellChat was performed on all T cells and *CD69*^+^*IGLL1*^+^ HSC-like pAML from RUNX1 samples with predominantly *CD69*^+^*IGLL1*^+^ HSC-like pAML. Cell subtypes with the strongest and most numerous interactions were inferred from CellChat’s outgoing/incoming interaction strength metrics and visualized using chord diagrams. The pathways underlying the most significant interactions were examined and visualized using heat maps.

*Identification of malignant mast cells in scRNAseq and bulk RNAseq*

Mast-like pAML cells were identified as HRTR subtypes in both RUNX1 and CBFB patients. To examine the role of mast cells in pAML we analyzed the scRNAseq dataset and TARGET bulk RNAseq dataset. Mast cells were identified in scRNAseq based on the expression of canonical markers: *TPSAB1, TPSB2, IL5RA*, and *FCER1A*. The proportion of mast cells in RUNX1 and CBFB (collectively the “core binding factor”, CBF pAMLs) was calculated by dividing the number of mast cells from a patient by the total number of detected cells for that patient. The proportions of mast cells in CBF patients were compared against all other cytogenetic categories using the student’s t-test. To identify samples enriched for mast cells in the TARGET dataset, we calculated the log_2_ mean z-score + 1 of the normalized mast cell gene expression (*TPSAB1, TPSB2, IL5RA,* and *FCER1A*). Mast gene enrichment was compared between patients with CBF mutation and patients from other cytogenetic backgrounds using the student’s t-test.

*Validating mast cells’ malignant annotation*

Two approaches were used to validate the malignant annotation of mast cells in scRNAseq. First, genes known to be over-expressed in RUNX1-RUNX1T1 (*RUNX1T1, POU4F1*) and CBFB-MYH11 (*DUSP6, TES*)^16,17^ mutated AML were compared between four groups of cells: mast cells from RUNX1/CBFB patients, all non-mast pAML clusters from RUNX1/CBFB patients, all pAML clusters from other cytogenetic backgrounds, and all non-AML cells (e.g. T cells, B cells). Differential expression between these groups was performed using the FindMarkers function in Seurat with Wilcoxon rank-sum significance testing. Second, differential expression was performed between mast cells and all non-malignant cells from CBF samples using the FindMarkers function with Wilcoxon significance testing. The rank-ordered gene list was used for gene set enrichment analysis in the ROSS_ACUTE_MYELOID_LEUKEMIA_CBF gene set, with the interpretation that if mast cells expressed genes enriched in this pathway it would indicate malignancy^17^. Systemic mastocytosis is a neoplasm of mast cells that can occur independently or in association with myeloid leukemia^18^. In >90% of cases, systemic mastocytosis is driven by a mutation in the KIT gene^19^. To determine if KIT mutation contributed to the mastocytosis observed in CBF pAML, we compared mast cell gene expression between samples with and without KIT mutation to determine if the KIT mutation was potentially driving the observed mast cell enrichment in CBF patients. The same mast enrichment score as used for the estimation of mast abundance was used to compare Kit mutated and Kit not mutated samples by the student’s t-test with a significance threshold of 0.05.

*Determining if mast cells were associated with shortened overall survival in CBF pAML*

To determine if enrichment for mast cells was associated with poor outcomes, we stratified CBF mutated pAML TARGET samples into mast-enriched or depleted samples based on enrichment scores of mast cell genes including TPSB2, TPSAB1, ILRA5, and FCER1A as well as CD69 which was found to be highly expressed in both subtypes of mast-like pAML. In scRNAseq, 7/24 patients had >5% mast cells (29.1% of patients). Five percent is the previously reported cut point for elevated mast cells in adult AML studies^20^. Samples in TARGET were considered mast-enriched if the log_2_ mean z-score + 1 of mast genes was 2 standard errors above the mean for all diagnostic bone marrow biopsies from CBF pAML patients to produce a similar proportion of samples labeled “mast enriched”. Overall survival was compared between mast-enriched and mast-depleted samples by the Cox proportional hazards model using the Survival package^8^.

*Differential expression and pathway analysis of mast cells against other pAML*

To understand the biological functions driving mast-like pAML in CBF patients we performed differential expression and pathway analysis. Differential expression was performed using the FindMarkers function with the Wilcoxon rank-sum test for significance testing between mast pAML and all other pAML subtypes. The differentially expressed genes (adjusted p-value < 0.05) were used for pathway analysis using the Reactome pathways in the EnrichR package^10,11^.
